# Peripheral and central employment of acid-sensing ion channels during early bilaterian evolution

**DOI:** 10.1101/2022.03.17.484724

**Authors:** Josep Martí-Solans, Aina Børve, Paul Bump, Andreas Hejnol, Timothy Lynagh

**Author notes:** Corresponding authors Timothy Lynagh, Sars Centre, UiB, Thormøhlensgate 55, 5008 Bergen, Norway, Andreas Hejnol, Institutt for Biovitenskap, UiB, Thormøhlensgate 53A, 5006 Bergen, Norway.

## Abstract

Nervous systems are endowed with rapid chemosensation and intercellular signaling by ligand-gated ion channels (LGICs). While a complex, bilaterally symmetrical nervous system is a major innovation of bilaterian animals, the employment of specific LGICs during early bilaterian evolution is poorly understood. We therefore questioned bilaterian animals’ employment of acid-sensing ion channels (ASICs), LGICs that mediate fast excitatory responses to decreases in extracellular pH in vertebrate neurons. Our phylogenetic analysis identified an earlier emergence of ASICs from the overarching DEG/ENaC superfamily than previously thought and suggests that ASICs were a bilaterian innovation. Our broad examination of ASIC gene expression and biophysical function in each major bilaterian lineage of Xenacoelomorpha, Protostomia, and Deuterostomia, suggests that the earliest bilaterian ASICs were probably expressed in the periphery, before being incorporated into the brain as it emerged independently in certain deuterostomes and xenacoelomorphs. The loss of certain peripheral cells from Ecdysozoa when they split from other protostomes likely explains their loss of ASICs, and thus the absence of ASICs in model organisms *Drosophila* and *C. elegans*. Thus, our use of diverse bilaterians in the investigation of LGIC expression and function offers a unique hypothesis on the employment of LGICs in early bilaterian evolution.

## Introduction

Morphological and behavioral complexity of animals is facilitated by nervous systems in which peripheral neurons sense and convey information from the environment, and more central neurons integrate and dispatch information to effectors (1, 2). Such sensory (environment → cell) and synaptic (cell → cell) signals rely on ligand-gated ion channels (LGICs), membrane proteins that convert chemical messages into transmembrane ionic currents within milliseconds (3, 4). LGICs are found in all multicellular animals (Metazoa) and even in outgroup lineages such as bacteria and plants (5-7). Compared to animals without nervous systems (Porifera and Placozoa), however, animals with nervous systems (Ctenophora, Cnidaria, and Bilateria) present a larger and more diverse LGIC gene content, i.e. more Cys-loop receptors, ionotropic glutamate receptors, and degenerin/epithelial sodium channel (DEG/ENaC) genes (8). Although the evolution of the original nervous system(s) did not necessarily involve a rapid expansion of LGICs, the elaboration and refinement of nervous systems within particular lineages did, and this likely endowed bilaterian neurons with a sophisticated chemo-electric toolbox in the ancestors of today’s complex bilaterian animals (1, 8, 9). Unfortunately, studies addressing the functional contribution of LGICs to early bilaterians are lacking (10), and we therefore lack crucial insight into evolution of the nervous system.

Diversity within the DEG/ENaC superfamily of channels exemplifies the novel tools that LGICs can offer an evolving nervous system. Several independent expansions of DEG/ENaC subfamilies have occurred, including degenerin channels (DEGs) in nematodes, pickpocket channels (PPKs) in arthropods, peptide-gated channels (HyNaCs) in cnidarians, and acid-sensing ion channels (ASICs) in vertebrates (11-15) (Fig. 1A). ASICs are excitatory sodium channels gated by increased proton concentrations (drops in pH) and are widely expressed in the nervous system of rodents, with scattered expression in other cells, such as epithelia (16) (Fig. 1B,C). ASICs in sensory neurons of skin, joints, and the gastrointestinal tract of rodents and humans contribute to pain, hyperalgesia, and touch (17-20), typifying a chemosensory role, whereas ASICs expressed postsynaptically in central neurons of rodents mediate depolarization in response to brief drops in synaptic pH during neurotransmission (21, 22), indicating an additional, central and synaptic role. ASICs occur in all deuterostomes (12), one of the three major lineages within Bilateria (Fig. 1B). But given the central or peripheral expression of ASICs in different deuterostomes (12, 23, 24) and uncertainty over the lineage in which ASICs emerged (Fig. 11B), we lack an accurate assessment of how ASICs were deployed by diversifying bilaterians and thus the chance of a better understanding of how complex bilaterian nervous systems evolved. This would require a broad and comprehensive assessment of ASICs throughout the major bilaterian lineages.

**Fig. 1.**
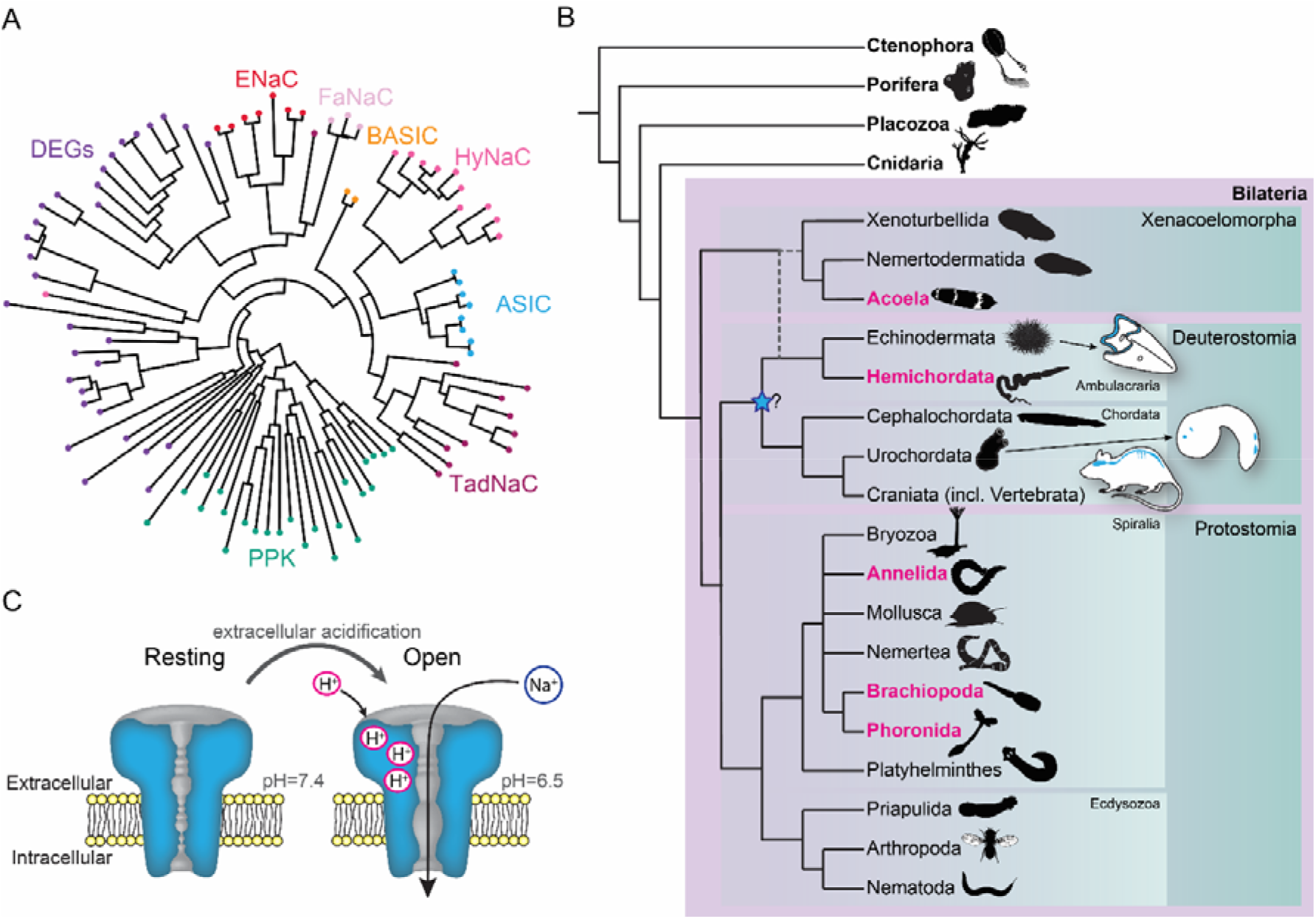
Overview of ASICs in Metazoa. (A) Abridged phylogenetic tree of the DEG/ENaC family showing major expansions of previously studied members (after (11, 25)). (B) Phylogenetic tree of main animal groups highlighting groups studied in this work (magenta), known ASIC expression (blue) and previously suggested ASIC emergence (blue star). Dotted lines show alternative positions for Xenacoelomorpha. Cartoons show previously described and well established ASIC protein and/or RNA expression in Chordata (rodent brain, spinal cord and sensory ganglia (26); urochordate larva sensory vesicle and bipolar tail neurons (23)) and Ambulacraria (echinoderm larva ciliary band (24)). (C) ASIC function. At rest, the channel is closed and impermeable. Upon extracellular acidification, certain amino acid side chains in the ASIC extracellular domain are protonated, causing conformational changes that open the channel, allowing sodium ions to flow down their chemo-electric gradient across the membrane.

We therefore performed a thorough phylogenetic investigation of metazoan DEG/ENaC genes, with a focus on ASICs, using unexplored transcriptomes and genomes, revealing ASICs throughout the Bilateria. Moreover, we analyzed gene expression using in situ hybridization and determined the electrophysiological properties of diverse ASICs from each major bilaterian lienage. Results from these experiments enable a new hypothesis on the evolution and function of ASICs during bilaterian evolution.

## Results

### Broader phylogenetic study identifies ASICs in Protostomia and Xenacoelomorpha

We first sought a definitive picture of how broadly ASIC genes are conserved throughout the five metazoan lineages of Bilateria, Cnidaria, Porifera, Placozoa, and Ctenophora (see Fig. 1B for phylogenetic relationship of these lineages). To this end we utilized previously unexplored transcriptomes combined with canonical resources to search for DEG/ENaC genes from all lineages. The phylogenetic analysis of these 700 sequences from 47 species shows a well-supported clade of ASICs (Fig. 2 and Fig. S1). The ASIC branch consists of two sub-clades “A” and “B”, both of which include bona fide ASICs from deuterostome bilaterians (12). No ASICs were identified in Cnidaria, Porifera, Placozoa, and Ctenophora.

**Fig. 2.**
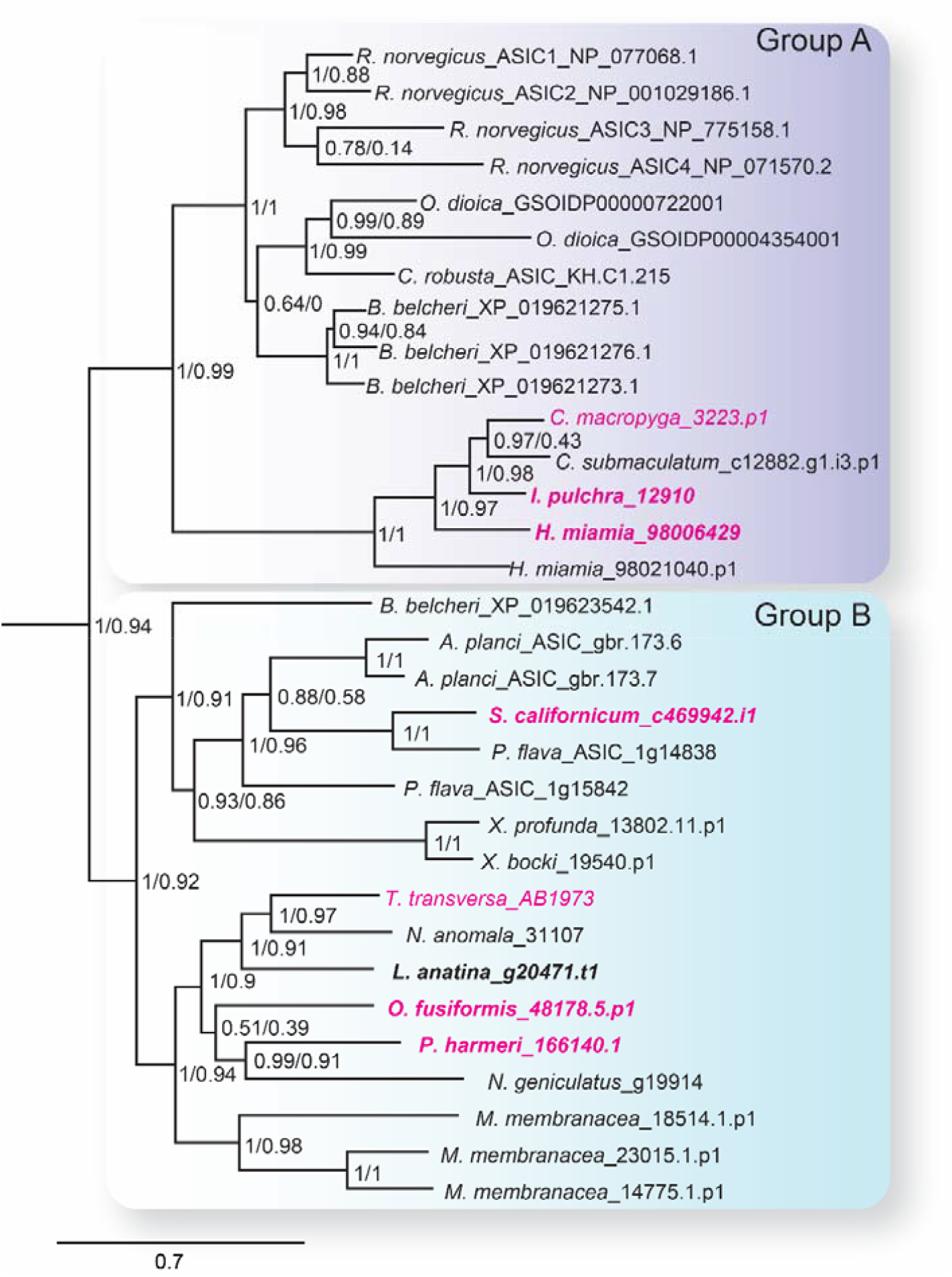
The ASIC branch of the DEG/ENaC family includes only bilaterian genes. ASIC branch from phylogenetic tree of DEG/ENaC family including 700 amino acid sequences from 47 metazoans (Fig. S1). Genes analyzed experimentally in this study are in magenta (gene expression) and/or bold (electrophysiology). Scale bar, amino acid substitutions per site. aBayes (left) and aLRT SH-like (right) likelihood-based support values indicated.

Bilateria is divided into three major groups, Deuterostomia (above), Protostomia (e.g. arthropods and molluscs), and Xenacoelomorpha (acoelomorph flatworms without specialized nephridia). In contrast to previous studies, we detected ASICs in protostomes and xenacoelomorphs (Fig. 2). These include putative ASICs from seven protostome species—the annelid *Owenia fusiformis*, the nemertean *Notospermus geniculatus*, the brachiopods *Terebratalia transversa, Novocrania anomala* and *Lingula anatina*, the phoronid *Phoronopsis harmeri*, the bryozoan *Membranipora membranacea*, and six xenacolomorph species, the acoels *Hofstenia miamia, Isodiametra pulchra, Convolutriloba macropyga*, and *Childia submaculatum*, and the xenoturbellans *Xenoturbella bocki* and *Xenoturbella profunda*. Our analysis also included the hemichordate *Schizocardium californicum*, whose ASIC groups with previously reported hemichordate and echinoderm ASICs (12), indicative of broad conservation of ASICs in Ambulacraria, the deuterostome group including hemichordates and echinoderms. This shows that ASIC genes are present in the three major bilaterian groups of deuterostomes, protostomes, and xenacoelomorphs and absent from all other lineages, suggesting that ASICs diverged from other DEG/ENaC genes after the Cnidaria/Bilateria split and before the Bilateria diversified.

Protostomes are divided into Spiralia (such as annelids, molluscs, brachiopods, and phoronids), and Ecdysozoa (such as the nematode *Caenorhabditis elegans* and arthropod *Drosophila melanogaster*) (Fig. 1B). Notably, the ASIC clade includes no ecdysozoan genes, although our analysis included DEG/ENaC genes from the nematode *Pontonema vulgare*, pan-arthropods *Centruroides sculpturatus* and *Daphnia pulex*, and priapulids *Priapulus caudatus* and *Halicryptus spinulosus*. We also see no putative ASICs in certain spiralians, including the molluscs *Acanthopleura granulata* and *Crassostrea gigas*, the platyhelminths *Prostheceraeus vittatus* and *Schmidtea mediterranea* and in certain xenacoelomorphs, including the nemertodermatids *Meara stichopi* and *Nemertoderma westbladi*. This analysis suggests that ASICs, although found throughout Bilateria, were lost in the lineage to the Ecdysozoa shortly after the Ecdysozoa/Spiralia split and were also lost in certain other bilaterian lineages such as molluscs and nemertodermatids.

### ASICs are expressed in two domains in Xenacoelomorpha

These results suggest that ASICs emerged during early bilaterian evolution, and we next sought evidence for potential biological roles by investigating the expression of ASIC genes throughout the major bilaterian lineages. The precise relationships between these lineages are under debate, but Xenacoelomorpha forms a putative sister group to all remaining Bilateria (27-30). Therefore, we investigated ASIC expression in Xenacoelomorpha, utilizing the acoels *Isodiametra pulchra, Hofstenia miamia*, and *Convolutriloba macropyga*. Xenacoelomorphs show a large variation in nervous system architecture, but all possess a basal epidermal nerve plexus and some species, more internally, an anterior condensation of neurons or “brain” (31). *I. pulchra* and *C. macropyga* also possess longitudinal bundles of neurons (32-34). External to this nervous system in acoels is generally a sheet of longitudinal and ring muscles and, more externally, ciliated epithelial cells mediating locomotion (35-37).

From a dorsal view, expression of *I. pulchra* ASIC mRNA can be detected in several cells across the central anterior of the adult animal (Fig. 3A). This is the location of the *I. pulchra* brain, where e.g. serotonergic and peptidergic neurons form lateral lobes, connected by a frontal ring and a posterior commissure, and from which four pairs of nerve cords extend posteriorly (Fig. 3A (32)). Fluorescent confocal micrographs show that the ASIC-expressing cells are associated with the posterior commissure of the brain and the dorsal and lateral neurite bundles (Fig. 3A, white arrowheads). Based on additional colorimetric *in situ* hybridization showing neuronal marker expression, the *I. pulchra* ASIC expression overlaps most closely with monoaminergic neurons of the brain (Fig. S2A). In *H. miamia*, we observed high ASIC expression in cells throughout the anterior third of the animal (Fig. 3B), correlating with the brain-like anterior condensation of neurons in this acoel (38). Similar to *I. pulchra*, the ASIC signal is strongest near the dorsal commissure but, in contrast, also in scattered cells across the animal, including in the very periphery (Fig. 3B, yellow arrowhead). *C. macropyga* ASIC expression was high and broad. In the anterior, ASIC-expressing cells are clearly associated with the brain (Fig. 3C’, white arrowheads). Throughout the animal, including the very periphery, there is medium to high ASIC expression, perhaps associated with locomotory ciliated cells covering the animal or the extensive nerve plexus (Fig. 3C’’). In summary, *I. pulchra* shows localized expression of ASIC in the brain, whereas *C. macropyga* and *H. miamia* ASIC show this central expression in addition to scattered, peripheral expression. Thus, ASIC is expressed in two distinct domains in Xenacoelomorpha, central and peripheral, and ASICs could have performed central, integrative, and peripheral, sensory roles in early acoels.

**Fig. 3.**
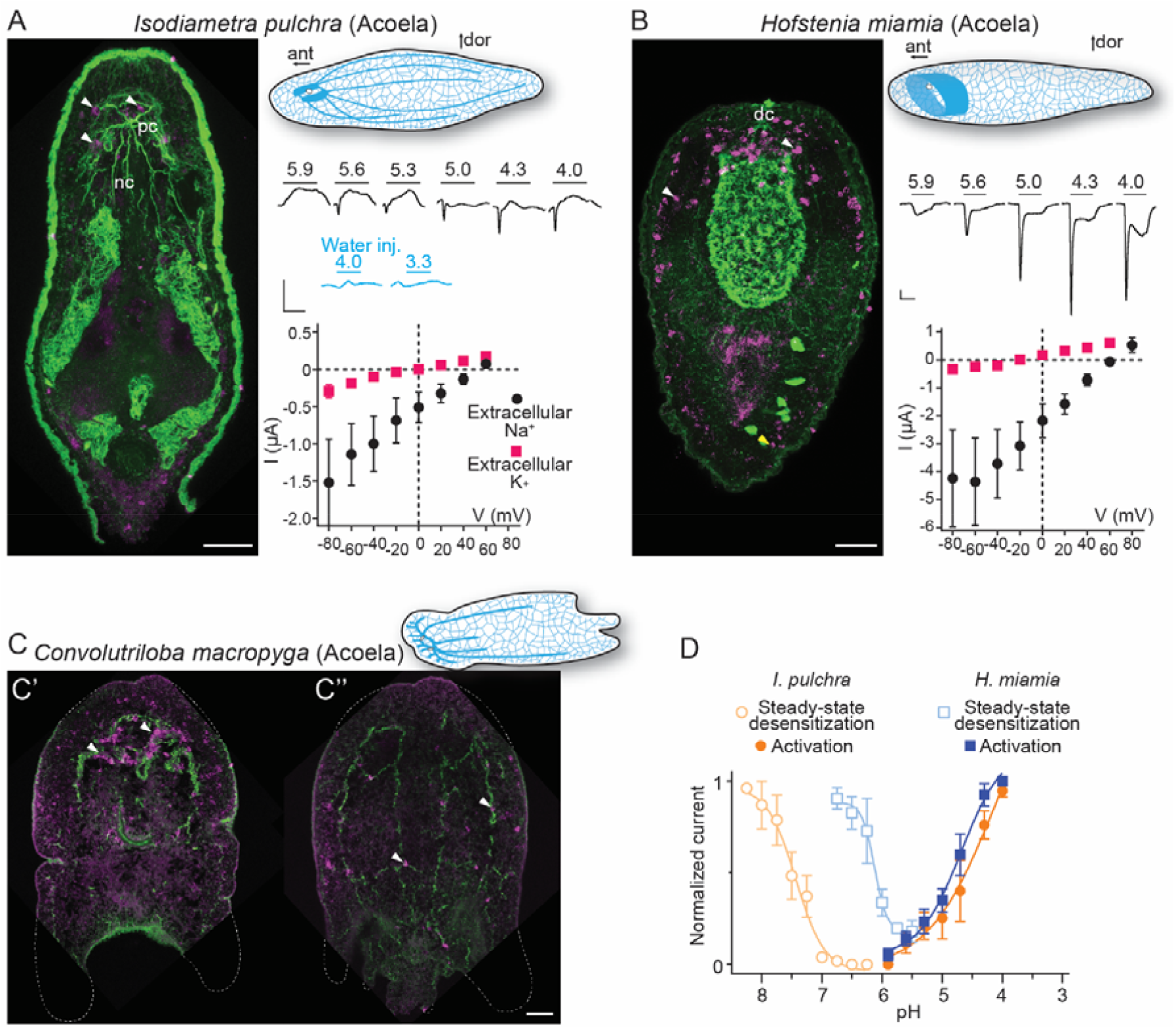
Expression and function of ASICs in Xenacoelomorpha. (A,B) Left: fluorescent confocal micrograph showing ASIC mRNA expression (magenta) and tyrosinated tubulin immunoreactivity (green), scale bar 40 µm. Upper right: cartoons illustrating morphology and nervous system (blue) after (33). ant, anterior; dor, dorsal; dc, dorsal commissure; nb, neurite bundles; pc, posterior commissure. Mid-right: proton-gated currents in *Xenopus laevis* oocytes expressing xenacoelomorph ASICs or water injected (scale bars: x, 5 s; y, 0.5 μA). Lower right: Mean (± SEM) pH 4-gated current (I, μA) at different membrane potentials (V, mV) in the presence of 96 mM extracellular NaCl or KCl (n = 7-8). Reversal potential (V_rev_) was read off these plots and the difference between V_rev,NaCl_ and V_rev,KCl_ was used to calculate relative ion permeability (P_Na+_/P_K+_, *Materials and Methods*). (C) ASIC mRNA expression (magenta) and tyrosinated tubulin immunoreactivity in *C. macropyga*. As *C. macropyga* is larger, images of slightly ventral (C’) and dorsal (C’’) planes are shown. (D) Filled symbols: mean (± SEM) normalized current amplitude in response to increasing proton concentrations (activation, n = 8). Open symbols: mean (± SEM) normalized current amplitude in response to pH 4 following pre-incubation in decreasing pH (steady state desensitization, n = 4-6).

### Xenacoelomorph ASICs mediate excitatory currents gated by high proton concentrations

When ASICs are exposed to drops in extracellular pH they typically show a transient depolarizing current (inward flow of Na^+^ ions), rapidly followed by either desensitization—a non-ion-conducting state in the presence of agonist—or a smaller sustained current (39). To test the function of xenacoelomorph ASICs, we injected the ASIC cRNAs into *Xenopus laevis* oocytes and measured membrane current in response to decreasing pH using two-electrode voltage clamp. At oocytes expressing *I. pulchra* ASIC, drops to pH 5.6 and lower rapidly activated a transient current (Fig. 3A, mid-right). However, compared to other ASICs, responses to increased proton concentrations were relatively inconsistent at *I. pulchra* ASIC: although transient currents were never activated by pH higher than 5.6, the concentration dependence of the transient current between pH 5.6 and 4.0 was inconsistent. To verify that these variable currents were indeed specific to the heterologous *I. pulchra* channels, we also applied pH 4.0 to oocytes from the same batch injected with water (Fig. 3A mid-right) or with a proton-insensitive channel (HyNaC2/4/6; Fig. S1), and we did not observe pH 4.0-gated currents (Fig. S1). Thus, *I. pulchra* ASIC is indeed gated by protons, but with relatively low potency (pH for half-maximal activation of transient current (pH_50_) = 4.0 ± 0.2). The *H. miamia* ASIC 98006429 also formed homomeric proton-gated channels, with rapid transient currents followed by smaller sustained currents in response to drops to pH 5.6 through 4.0 (pH_50_ = 4.8 ± 0.1, Fig. 3B,D). The presence of another ASIC gene in this species, 9021040 in Fig. 2, raises the possibility of heteromeric channels in Xenacoelomorpha, but we have not pursued that here. Finally, we observed no proton-gated currents in *Xenopus* oocytes injected with *C. macropyga* ASIC cRNA (n = 6), suggesting either low heterologous expression or proton insensitivity of this channel. Like vertebrate ASICs, *I. pulchra* and *H. miamia* ASICs showed decreased responses to activating pH (4.0) after preincubation in slightly decreased pH (Fig. 3D, open symbols), indicating that steady-state desensitization is a broadly conserved phenomenon.

Most ASICs so far studied have a slight preference for sodium over potassium ions, with a relative sodium/potassium ion permeability (P_Na+_/P_K+_) of ∼10, and thus activation of ASICs leads to depolarization and generation of action potentials in mice (40). *I. pulchra* ASIC (P_Na+_/P_K+_ = 8.3 ± 2.0) showed similar Na^+^ selectivity to most ASICs (Fig. 3A, lower right), whereas *H. miamia* Na^+^ selectivity (P_Na+_/P_K+_ = 36 ± 11; Fig. 3B) is remarkably high compared to most ASICs and is more reminiscent of the high preference for sodium seen in ENaCs (41). Thus, both in xenacoelomorphs, which belong to the putative sister group to all other bilaterians, and in vertebrates such as rodents and humans, ASICs are excitatory receptors for protons that are expressed highly in both the brain and in the periphery.

### ASICs are expressed in peripheral neurons and the digestive system in Spiralia

Excitatory proton-gated currents and combined central and peripheral expression of xenacoelomorph ASICs is reminiscent of vertebrate ASICs. However, whether the ancestral bilaterian ASIC performed similar roles remains unclear, as the position of Xenacoelomorpha as sister taxon of all other bilaterians or as part of a group with Ambulacraria within Deuterostomia is debated (27-30)(Fig.1B). We therefore turned to the third major lineage of bilaterians and investigated previously unidentified ASICs in Protostomia. We investigated the expression and function of ASICs in brachiopod, phoronid, and annelid larvae, allowing us to study whole-animal gene expression in animals with typical spiralian features such as an anterior brain or apical organ and ventrolateral or peripherally extending nerve cords.

In the brachiopod *Terebratalia transversa*, ASIC was expressed most highly in cells at the lateral edge of the central neuropil in late larvae (Fig. 4A). These cells are slightly lateral/caudal to previously identified sensory neurons in the *T. transversa* apical organ (42), and additional *in situ* hybridization targeting distinct neuronal types showed that the ASIC-expressing cells are close to or overlap with cholinergic neurons at the lateral edge of the central neuropil (Fig. S2B). Confocal images suggest that these ASIC-expressing cells project latero-anteriorly (Fig. 4A, inset), perhaps indicative of an anterior sensory role. In the phoronid *Phoronopsis harmeri*, ASIC was expressed in two principal domains. There was a clear signal in relatively few cells in the perimeter of the hood (Fig. 4B, black arrowheads), a domain innervated by the peripheral nerve ring and controlling swimming by ciliary beating (43, 44). There was no ASIC expression in the more central parts of the nervous system, including both the apical organ, an anterior group of sensory neurons that gives way to the developing brain, and the main nerve ring that runs caudally from the apical organ through the trunk and innervates the tentacles (44-46). The second domain, in which expression was even more prominent, was the digestive system. Here, expression was high in numerous cells around the digestive tract (comprising a stomach in the trunk and intestine in the pedicle at this stage), most noticeably just below the pharynx and around the lower half of the stomach (Fig. 4B, orange arrowheads, (47)). We also detected digestive system expression in the more distantly related spiralian, the annelid *Owenia fusiformis*. In 19 hours post fertilization *O. fusiformis*, we observed no ASIC expression in the young nervous system consisting dorsal apical organ, ventral prototroch (or ciliary band) neurons, and anterior and lateral connecting neurons (48, 49). Instead, ASIC was clearly expressed in cells around the oesophagus, and, more broadly, in the midgut (Fig. 4C). These results show that in various spiralians, ASICs are found in the periphery and the digestive system, not in the brain (or apical organ).

**Fig. 4.**
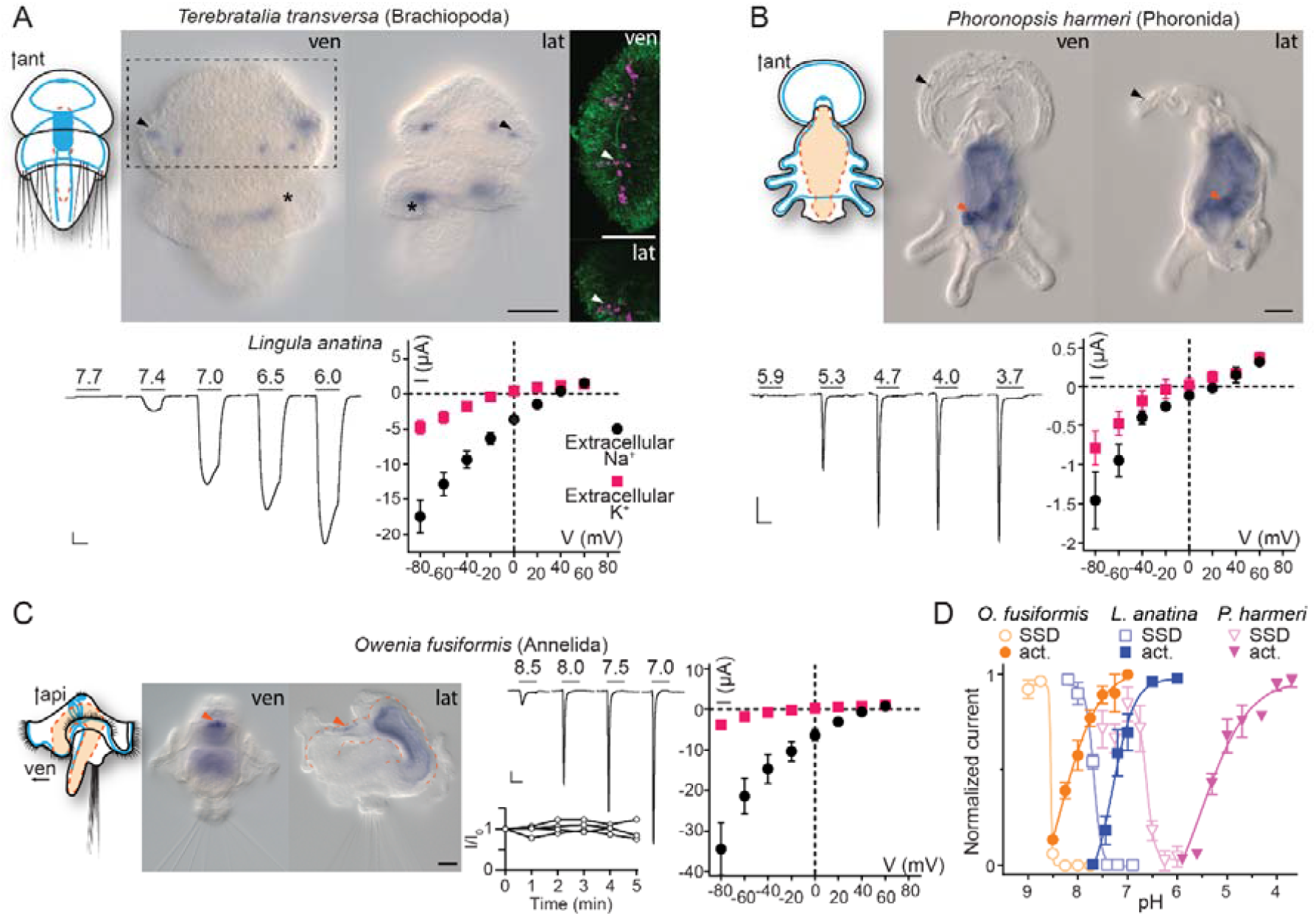
Expression and function of ASICs in Spiralia. (A,B) Upper: cartoons illustrating nervous system (blue) and digestive system (orange; Anlage only in *T. transversa*) after (49, 50) and ASIC mRNA expression (ant, anterior; api, apical; lat, lateral; ven, ventral; scale bar, 40 µm). Arrowheads, ASIC expression; asterisks, unspecific staining common in *T. transversa* (33). Lower left: proton-gated currents in *Xenopus laevis* oocytes expressing indicated spiralian ASICs (scale bars: x, 5 s; y, 1 μA). Lower right: mean (± SEM) pH 6.5- (*L. anatina*) or 4- (*P. harmeri* ASIC) gated current (I, μA) at different membrane potentials (V, mV) in the presence of 96 mM extracellular NaCl or KCl (n = 5-8). Reversal potential (V_rev_) was read off these plots and the difference between V_rev,NaCl_ and V_rev,KCl_ was used to calculate relative ion permeability (P_Na+_/P_K+_). (C) Left to right: Animal morphology, ASIC expression, proton-gated currents (top) and normalized current amplitude, i.e. amplitude of the current at certain time (I) divided by the current amplitude at the start of the experiment (I_0_), in response to same proton concentration (pH 7.0) in oocytes perfused with pH 9.0 solution for 5 minutes (bottom), lines connect data from individual oocytes, and pH 6.5-gated current at different membrane potentials, as in (A,B) at *O. fusiformis* ASIC. (D) Filled symbols: mean (± SEM) normalized current amplitude in response to increasing proton concentrations (activation, “act.”, n = 6-11). Open symbols: mean (± SEM) normalized current amplitude in response to pH 7 for *O. fusiformis*, 6.5 for *L. anatina*, and 4 for *P. harmeri* ASIC following pre-incubation in decreasing pH (steady state desensitization, n = 4-5).

### Spiralian ASICs show a wide range of proton sensitivity and ion selectivity

We next tested the electrophysiological properties of spiralian ASICs by expressing them heterologously in *Xenopus* oocytes. We observed no proton-activated currents in *Xenopus* oocytes injected with *T. transversa* (brachiopod) ASIC RNA (n = 10), and we cannot conclude if this is due to low heterologous expression or unknown function of this channel. However, we identified an ASIC transcript in another brachiopod, *Lingula anatina* (51), and observed functional expression of this channel. *L. anatina* ASIC was sensitive to relatively low proton concentrations, activated by pH in the range of 7.4 to 6.0 (Fig. 4A, pH_50_ = 7.3 ± 0.2) and showed typical selectivity for sodium over potassium ions (Fig. 4A, P_Na+_/P_K+_ = 9.0 ± 1.7). The phoronid *P. harmeri* ASIC showed much lower proton sensitivity, with small currents in response to pH 5.3 and lower that rapidly desensitized in the continued presence of low pH (Fig. 4B, pH_50_ = 5.1 ± 0.1) with almost no preference for sodium over potassium ions (Fig. 4B, P_Na+_/P_K+_ = 3.0 ± 0.9). We next tested the function of ASIC from the annelid *O. fusiformis*. This channel was also highly sensitive to low proton concentrations. When held at pH 7.5 and exposed to lower pH, *O. fusiformis* ASIC showed very small current responses, but when held at pH 9.0 and exposed to small drops in pH, large inward currents were activated that rapidly desensitized (Fig. 4C, pH_50_ = 8.1 ± 0.1). We measured repeated responses of *O. fusiformis* ASIC to pH 7.0 over the course of five minutes in pH 9.0, and observed no decrease in current amplitude (Fig. 4C), verifying that the relatively basic resting pH of 9.0 was not harming oocytes and skewing results with this channel. *O. fusiformis* ASIC also showed ion selectivity typical of most ASICs (Fig. 4C, P_Na+_/P_K+_ = 12.0 ± 0.3).

Spiralian ASICs seem to vary in apparent proton affinity and in ion selectivity among the lineages, from low proton sensitivity (pH ∼5) and essentially non-selective cation currents in phoronid ASIC to high proton sensitivity (pH 7-8) and ∼10-fold selective sodium permeability in brachiopod and annelid ASICs. All showed canonical steady-state desensitization, occurring in pH ranges higher than activating pH (Fig. 4D). Spiralian ASIC genes thus encode proton-gated cation channels, and ASICs are present in all major groups of bilaterians: Xenacoelomorpha, Protostomia, and Deuterostomia.

### Hemichordate ASIC is expressed in peripheral cells and pharynx and mediates rapidly desensitizing excitatory currents in response to protons

Regarding Deuterostomia, combined expression and function of ASICs have so far only been characterized in selected chordates (urochordates and vertebrates) (12, 23, 52), and relatively little is known about ASICs in the ambulacrarian lineage (echinoderms and hemichordates). Recent studies on neural development in sea urchin (*Lytechinus variegatus*, an echinoderm) larvae showed that ASIC is expressed diffusely throughout the ciliary band (24). This transcript (MH996684) is closely related to the Group B ASICs from echinoderms and hemichordates that showed proton-gated currents when expressed heterologously (12). Here, we used the hemichordate *Schizocardium californicum* to characterize the combined expression and function of an ambulacrarian ASIC. *S. californicum* ASIC expression was visible in late larval stages when structures such as the nervous system and the ciliary bands become more intricate, like sea urchin ASIC (24). The tornaria larva of *S. californicum* has two main ciliary bands: the circumoral band for feeding; and the telotroch, innervated by serotonergic neurites, for locomotion (53). *S. californicum* ASIC expression was detected in both ciliary bands (Fig. 5A, black arrowheads) as well as in the pharynx (Fig. 5A, orange arrowhead), however, signal intensity was variable between specimens and sometimes difficult to detect in all domains. When heterologously expressed, *S. californicum* ASIC showed rapidly activating and desensitizing currents in response to extracellular acidification with proton sensitivity comparable to other hemichordate ASICs but lower than echinoderm ASICs (Fig. 5B, pH_50_ = 5.3 ± 0.1) (12). The *S. californicum* channel showed relatively weak ion selectivity, with P_Na+_/P_K+_ values slightly above unity (2.9 ± 0.5; Fig. 5D), similar to ion selectivity in previously described ambulacrarian ASICs (12). *S. californicum* thus presents a functional ASIC that is expressed in the peripheral nervous system and the digestive system, and it appears that such peripheral expression of ASICs is a conserved feature of Deuterostomia, Protostomia, and Xenacoelomorpha.

**Fig. 5.**
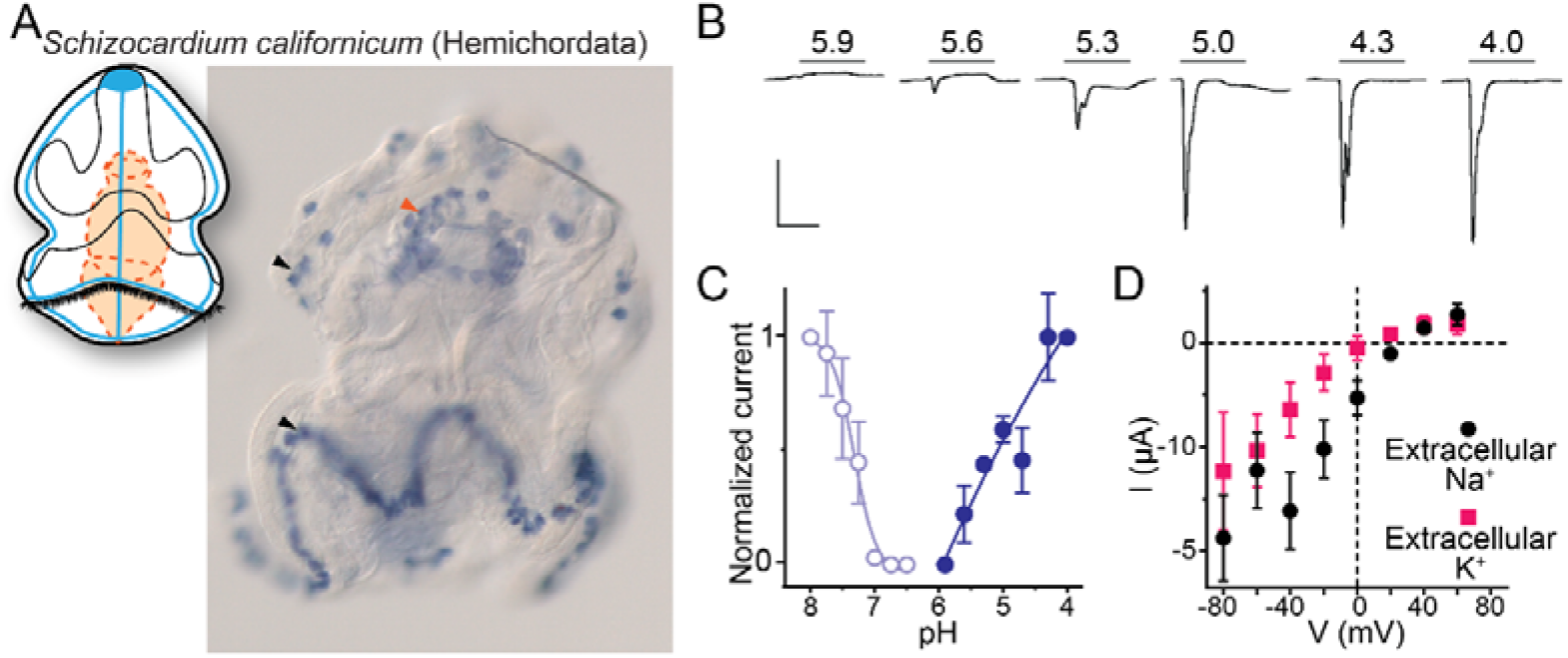
Expression and function of hemichordate ASIC. (A) Cartoon illustrating nervous system (blue) and digestive system (orange) in *Schizocardium californicum*, after (53), and colorimetric in situ hybridization showing *S. californicum* ASIC expression. Scale bar: 50 μm. (B) Proton-gated currents in *Xenopus laevis* oocytes expressing *S. californicum* ASIC. Scale bars: x, 5 s; y, 2 μA. (C) Filled symbols: mean (± SEM) normalized current amplitude in response to increasing proton concentrations (activation, “act.”, n = 5). Open symbols: mean (± SEM) normalized current amplitude in response to pH 4 following pre-incubation in decreasing pH (steady state desensitization, n = 4).

## Discussion

### The emergence of ASICs

Our results show that ASICs – of the same family as the prototypical rat ASIC1a – are present in Xenacoelomorpha and Protostomia in addition to Deuterostomia and are thus conserved in the three major groups of Bilateria. We also find, consistent with previous studies, that ASICs of this family are absent from the other major lineages of Cnidaria, Placozoa, Porifera, and Ctenophora, indicating that ASICs emerged in the lineage to the Bilateria, soon after the Cnidaria/Bilateria split ∼680 Mya (54). The clades most closely related to ASICs within the DEG/ENaC superfamily tree include three from which several channels have been characterized: mammalian bile acid-sensitive ion channels (BASICs), *Trichoplax adhaerans* Na^+^ channels (TadNaCs); and *Hydra vulgaris* and *Nematostella vectensis* Na^+^ channels (HyNaC and NeNaCs). We cannot establish phylogenetically the identity of the ancestral gene from which ASICs emerged, as branch support toward the base of the ASIC+BASIC+TadNaC+HyNaC/NeNaC clade is relatively low: our maximum likelihood trees inferred with aLRT SH-like and aBayes statistics yielded slightly different topologies within this clade (Fig. S1C,D), and previous studies using similar methods to each other also inferred slightly different topologies within this clade (55, 56). Statistical support for the distinct ASIC clade is strong, however (Fig. S1).

We are thus left to consider the *functional* relationships among these cousins. BASICs, highly expressed in mammal intestines, are activated by bile acids or are constitutively active, and show inhibition by protons (57, 58). Non-mammalian genes from the BASIC clade have not been characterized. HyNaCs are neuropeptide-gated channels from the medusozoan cnidarian *Hydra* (11), and when NeNaCs from the anthozoan cnidarian *Nematostella* were recently reported, surprisingly, NeNaC2 and NeNaC14 were activated by protons (pH_50_ values of 5.8 and <4.0), not by cnidarian neuropeptides (55). The TadNaC clade also includes a proton-activated channel, TadNaC2 (pH_50_ 5.1), in addition to a proton-inhibited channel, TadNaC6 (25, 56). Thus, proton-activated and -inhibited channels occur sporadically throughout the ASIC+BASIC+TadNaC+HyNaC/NeNaC clade, and only in the ASIC clade is proton-induced activation the defining feature. ASICs could thus have emerged from a proton-activated ancestor, after which this function was selected for; or from a peptide- or bile acid-activated ancestor, in which a novel mechanism of activation evolved. Certain functional and phylogenetic data have led others to favor the latter possibility (56, 59), and indeed structural features important for channel gating throughout the ASIC family, such as highly conserved histidine residues and/or aromatic interactions, are absent from proton-gated NeNaCs and TadNaCs (12, 56). Nonetheless, the fact that proton-induced activation has emerged in numerous DEG/ENaC channels, even outside the ASIC+BASIC+TadNaC+HyNaC/NeNaC clade, as shown for a *Drosophila* PPK channel and three *C. elegans* degenerin-like channels (60, 61), suggests a propensity for this function in the DEG/ENaC superfamily.

Soon after its emergence in an early bilaterian, we infer that ASIC duplicated, giving rise to two similar proton-gated channels, based on Group A and Group B ASICs in our phylogeny. Subsequently, and at different times, most descendants lost one of these (Fig. 6). Acoels lost Group B ASIC and xenoturbellans lost Group A ASIC soon after this early split within the Xenacoelomorpha. In contrast, the first deuterostomes and, even more recently, the first chordates likely retained both ASICs, reflected in the continued presence of both in Cephalochordata (Fig. 2, Fig. 6). Subsequently, in Olfactores (Craniata + Urochordata), Group B ASIC was lost and Group A ASIC underwent independent radiations, leading to e.g. ASIC1-ASIC4 paralogues in vertebrates (62), two Group A ASICs in urochordates (12), and multiple Group A ASICs in cephalochordates (Fig. 2).

**Figure 6.**
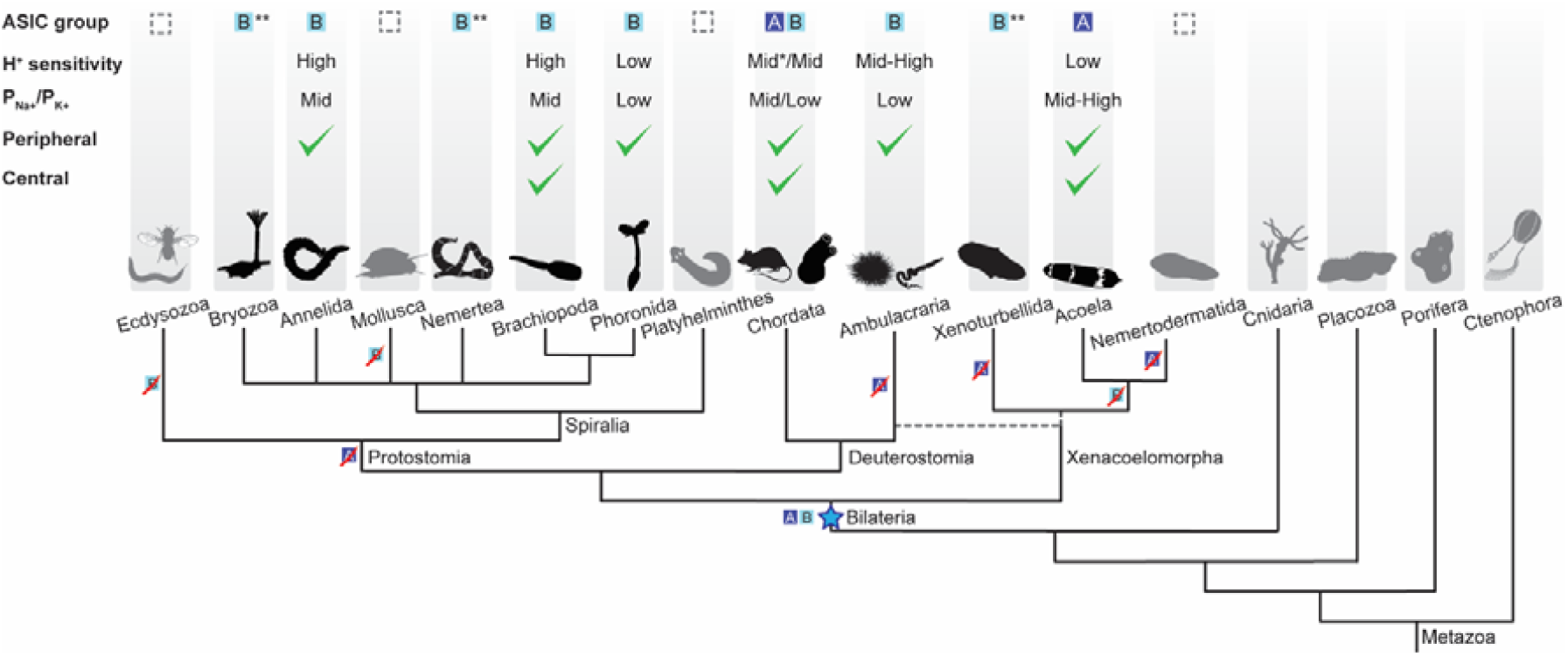
Evolutionary history of ASIC function in Metazoa. Upper half of diagram indicates characteristic properties of ASIC in different lineages: DEG/ENaC gene tree position (“ASIC group A or B”); proton (H^+^) sensitivity; relative ion permeability (“P_Na+_/P_K+_”); native expression pattern (“Peripheral”, ciliary or gastrointestinal epithelia and/or peripheral neurons; “Central”, brain and/or nerve cords). Lower half shows phylogenetic relationships of the different animal phyla studied here. Blue star, putative emergence of ASICs in the last common ancestor of all bilaterians after Cnidaria/Bilateria split. For H^+^ sensitivity, “low”, “mid”, and “high” correspond to pH_50_ <5.3, 5.3-7, and >7, respectively. *, a minority of characterized chordate ASICs have low H^+^ sensitivity ((41, 62)). For relative ion permeability, “low”, “mid”, and “high” correspond to P_Na+_/P_K+_ ≤3, 3-15, and >15, respectively. **, channel function not experimentally tested.

### Peripheral and central roles for ASICs

By investigating ASIC gene expression in each of the three major bilaterian lineages, we see that ASICs are principally found: centrally, in the brain; and/or peripherally, in the digestive system, peripheral nerve rings, and/or scattered peripheral cells. This, together with previous research on ASICs in rodents, points to at least two functions of ASICs: a central signaling role, where ASICs mediate rapid excitatory signals between neurons in the brain (21); and a peripheral sensory/modulatory role, where ASICs convert increased proton concentrations in the environment— extra-organismal or in the gut—into excitatory neuronal signals (63). Brain expression of mouse, zebrafish and urochordate ASICs suggests conservation of the synaptic role in various chordates (12, 21, 22, 62), whereas in the other deuterostome lineage of Ambulacraria (Fig. 1B), nervous systems are less centralized and ASIC expression appears more peripheral ((24) and Fig. 6). In Xenacoelomorpha, the brain expression pattern of ASIC resembles that of monoaminergic neurons and also overlaps with synaptotagmin expression (Fig. S2A). In contrast, central ASICs were observed in only one of the three Spiralia tested, and even these ASIC-expressing cells showed projections towards the very anterior of the animal, perhaps indicative of a less integrative and more sensory role (Fig. 4A). Brain ASICs are thus common to Xenacoelomorpha and Deuterostomia and are less prevalent in Protostomia, and this central role is more commonly played by Group A ASICs than Group B ASICs (Fig. 6).

The second, peripheral domain, observed in six of the seven species of Xenacoelomorpha, Spiralia, and Deuterostomia studied here, largely corresponds to the ciliation in the species. While xenacoelomorphs are completely covered with ciliated epithelial cells, in many other Bilateria the ciliation is restricted to ciliary bands or the ciliary ventral region, both used for locomotion. These ASICs are thus expressed in a location to sense external changes in pH and directly modulate ciliary movement via excitatory current or, via depolarization and synaptic release onto more central neurons, send the information centrally. Conversely, if these ASICs are expressed on the proximal side of these peripheral cells, their role may be modulation of ciliary function in response to efferent modulatory signals. Although activation of postsynaptic ASICs has so far been linked to only glutamatergic and GABAergic synapses (21, 64) and innervation of ciliary cells appears monoaminergic in Spiralia (43), monoaminergic vesicles are also acidic and could thus foreseeably release protons to activate ASICs here (65).

We were surprised to see expression of ASIC predominantly in the digestive system of the spiralians *O. fusiformis* and *P. harmeri*. In mammals, chemosensors such as ASIC and transient receptor potential channels (e.g. TRPV1) are expressed in sensory neurons innervating the gastrointestinal tract (17), but oesophageal, intestinal, and lung epithelial cells also express ASICs, potentially contributing to acid-induced secretions, transport, and inflammation (66-68). The gastrointestinal expression of ASICs that we observed may reflect a chemosensory role, like that mediated by ASICs in neurons innervating the lungs and skin of mammals (69, 70). Similar functions are tentatively suggested by ASIC expression in the peripheral nervous system of Ambulacraria ((24, 71), Fig. 5). Again, we can’t exclude the possibility that these peripheral ASICs are modulating ciliary function, either directly via acid-induced activation or indirectly, via efferent signals from central neurons. Nonetheless, these expression patterns seem to suggest that ASICs play a sensory role in the periphery of Deuterostomia, Protostomia, and Xenacoelomorpha via expression in cells that correlate with ciliation, indicating that such a role was probably present in the ancestral bilaterian. Consistent with the early appearance of this role, it is played by both Group A and Group B ASICs, which both emerged in early bilaterians. Subsequently, ASICs - particularly Group A - were likely deployed to the other, more central role for inter-neuronal communication in lineages such as Acoela and Chordata.

### Loss of ASICs

The conservation and high expression of ASICs in bilaterians begs the question as to why they were lost in certain lineages. The reason for the loss of ASICs in selected Spiralia, such as molluscs and platyhelminthes is unclear, especially when one considers that numerous molluscs are susceptible to the tide and thus vast fluctuations in pH. Presumably, gene radiations that increased sensory ion channel or transporter diversity in such animals have compensated for the loss of ASICs (72-76), and indeed, certain members of various channel families are capable of mediating excitation (or inhibition) in response to decreased pH (3). Perhaps most noticeable, however, is the absence of ASICs in Ecdysozoa, a broad lineage including pan-arthropods, nematodes, and priapulids, each of which we considered in our phylogenetic analysis. Ecdysozoa split from Spiralia ∼650 Mya (54), adopting a chitinous cuticle utilizing rigid locomotory structures and requiring periodic molting for growth (77, 78). Concomitantly, Ecdysozoa lost the ectodermal, motile ciliated cells inherited from the last common ancestor of Cnidaria and Bilateria that mediate locomotory, feeding, secretory, and sensory functions in most other bilaterians (79, 80). The correlation between loss of ASICs and loss of motile ciliated cells, together with the conservation of other ciliated and sensory cells in Ecdysozoa, indicates that the role of ASICs in early Protostomia was primarily associated with ectodermal ciliated cells rather than more central sensory cells associated with e.g. mouth, eyes, and antennae. This, together with the presence of ASICs in the periphery of Spiralia, Xenacoelomorpha, and Deuterostomia, suggests that the role of the earliest ASIC was likely local conversion of external chemical stimuli into modulation of locomotion, conversion of central signals into modulation of locomotion, or both. Indeed, motile ciliated epithelia in human lungs are modulated by both external and central stimuli (81), although the precise contribution of ASICs in these cells is unclear (67).

### Outlook

We acknowledge that future experiments might benefit from greater genomic resources for non-bilaterian animals, but our comprehensive survey of DEG/ENaC genes from Ctenophora, Porifera, Placozoa, and Cnidaria finds no ASICs in those lineages. Our study suggests that ASICs emerged in an early bilaterian, most likely in peripheral cells or epithelia, and were gradually adopted into the central nervous system of certain complex animals. This offers a unique insight into the employment of LGICs during early bilaterian evolution. Functional characterization of diverse ASICs shows a considerable breadth of pH sensitivity and ion selectivity throughout the family, offering new tools for probing the biophysical mechanisms of function. The combined use of gene expression and experimental analysis is thus a useful tool in understanding protein evolution and function.

## Materials and Methods

### Survey and phylogenetic analysis

Mouse ASIC1a was used as a query in tBLASTn searches of DEG/ENaC genes in xenacoelomorphs (*Convolutriloba macropyga, Childia submaculatum, Hofstenia miamia, Isodiametra pulchra, Xenoturbella bocki, Xenoturbella profunda, Nemertoderma westbladi*, and *Meara stichopi* from transcriptomes published in (82)), spiralians (*Spadella spp*., *Dimorphilus gyrociliatus, Epiphanes senta, Lepidodermella squamata, Lineus longissimus, Lineus ruber, Membranipora membranacea, Novocrania anomala, Owenia fusiformis, Phoronopsis harmeri, Prostheceraeus vittatus*, and *Terebratalia transversa* from our transcriptomes in preparation; *Acanthopleura granulata, Crassostrea gigas*, and *Brachionus plicatilis* from NCBI; *Lingula anatina* and *Notospermus geniculatus* from OIST; and *Schmidtea mediterranea* from SmedGD), ecdysozoans (*Halicryptus spinulosus, Priapulus caudatus, Pontonema vulgare* from our transcriptomes; *Drosophila melanogaster, Daphnia pulex, Centruroides sculpturatus* and *Caenorhabditis elegans* from NCBI) and a hemichordate (*Schizocardium californicum*, our transcriptome). Other DEG/ENaC genes were retrieved via BlastP at public databases NCBI, Compagen, JGI, OIST, OikoBase, Aniseed, or UniProt targeting cnidarians (hexacorallian *Nematostella vectensis*, octacorallian *Dendronephthya gigantea*, scyphozoan *Aurelia aurita*, hydrozoan *Hydra vulgaris*), poriferan (*Amphimedon queenslandica*), the placazoan *Trichoplax adhaerens*, ctenophores (cydippid *Pleurobrachia bachei* and lobate *Mnemiopsis leidyi*), and deuterostomes (chordates *Rattus norvegicus, Ciona robusta, Oikopleura dioica* and *Branchiostoma belcheri*; hemichordate *Ptychodera flava*; and echinoderm *Acanthaster planci*). Amino acid sequences were aligned using MAFFT (83), variable N- and C-termini were removed and highly similar sequences were not considered (Dataset S1), and homologies were assigned by phylogenetic tree analyses based on Maximum Likelihood (ML) inferences calculated with PhyML v3.0 (84). Robustness of tree topologies was assessed under automatic model selection based on Akaike Information Criteria. Due to computational load of bootstrap performance, trees were inferred using the fast likelihood-based methods: aLRT SH-like; and aBayes (85).

### Animal collection and fixation

Stable cultures of acoels were maintained in the laboratory. *Convolutriloba macropyga* (86) were reared in a tropical aquarium system with salinity 34+/-1 ppt at a constant temperature of 25 °C. The aquariums were illuminated (Pacific LED lamp WT470C LED64S/840 PSU WB L1600, Philips) on a day/night cycle of 12/12 hours. The worms were fed with freshly hatched brine shrimp *Artemia* twice per week. *Isodiametra pulchra* (87) were cultured as described by (88), and *Hofstenia miamia* (89) as described by (90). For the remaining species, adult gravid animals were collected from Bodega Bay, California, USA (*Phoronopsis harmeri* (91)), San Juan Island, Washington, USA (*Terebratalia transversa* (92)), Station Biologique de Roscoff, France (*Owenia fusiformis* (93)), and Kanangra Boyd National Park and Morro Bay State Park, California, USA (*Schizocardium californicum* (94)). Animals were spawned and larvae obtained as described in (95-98). Adult and larval specimens were starved for 2 - 7 days prior to fixation. Samples were relaxed in 7.4% magnesium chloride and fixed in 4% paraformaldehyde in culture medium for 1 hour at room temperature and washed several times in 0.1% Tween 20 phosphate buffered saline (PBS), dehydrated through a graded series of methanol, and stored in pure methanol or ethanol at −20 °C.

### Cloning

The full-length coding sequences of identified ASIC genes were amplified from cDNA of *C. macropyga, I. pulchra, H. miamia, P. harmeri, O. fusiformis, T. transversa*, and *S. californicum* by PCR using gene-specific primers. PCR products were purified and cloned into a pGEM-T Easy vector (Promega, A1360) according to the manufacturer’s instructions and the identity of inserts confirmed by sequencing. Riboprobes were synthesized with Ambion Megascript T7 (AM1334) and SP6 (AM1330) kit following manufacturer’s instruction for subsequent in situ hybridization.

### Immunohistochemistry and situ hybridization

Single whole-mount colorimetric and fluorescent in situ hybridization was performed following an established protocol (31) with probe concentration of 0.1 ng/μl (*I. pulchra*) or 1 ng/μl (the remaining species) and hybridization temperature of 67 °C. Proteinase K treatment time was adjusted for each species and ranged from 2 minutes (*P. harmeri, O. fusiformis, S. californicum*) to 10 minutes (*T. transversa)*. Post-hybridization low salt washes were performed with 0.05x saline sodium citrate (SSC; *H. miamia*) or 0.2x SSC (the remaining species). Fluorescent in situ hybridization was visualized with TSA Cy3 kit (PerkinElmer, NEL752001KT). Samples were mounted in 70% glycerol or subjected to immunohistochemistry for visualization of neural structures: samples were permeabilized in 0,2 % Triton-x in PBS (PTx) and blocked in 1% bovine serum albumin in PTx (PBT) and incubated with antibodies against tyrosinated tubulin (Sigma, T9028) at a concentration of 1:250 in PTx with 5% normal goat serum, and incubated for 16-18 hours at 4 °C. After several washes in PBT the samples were incubated with secondary goat anti-mouse antibodies conjugated with AlexaFluor 488 (Life Technologies), at a concentration 1:200 in PTx with 5% normal goat serum for 16-18 hours at 4 °C, and samples washed extensively before mounting in 70 % glycerol and imaging. Nuclei were stained with DAPI (Molecular Probes).

### Imaging

Representative specimens from colorimetric in situ hybridization experiments were imaged with a Zeiss Axiocam 503 color connected to a Zeiss Axioscope 5 using bright-field Nomarski optics. Fluorescently labelled samples were scanned in an Olympus FV3000 confocal laser-scanning microscope. Colorimetric in situ’s stained with antibodies were scanned in a Leica SP5 confocal laser-scanning microscope using reflection microscopy protocol as described by (99). Images were analyzed with Imaris 9.8.0 and Photoshop CS6 (Adobe), and figure plates were assembled with Illustrator CC. Brightness/contrast and color balance adjustments were applied to the whole image, not parts.

### Electrophysiological recordings and data analysis

For expression in *Xenopus laevis* oocytes and electrophysiological experiments, coding sequences were mutated synonymously to remove internal restriction sites if necessary and subcloned into SalI and XbaI sites of a modified pSP64poly(A) vector (ProMega), containing 5’ SP6 sequence, 5′- and 3′-UTR sequences of the *X. laevis* β-globin gene, and a C-terminal Myc tag, with an EcoRI restriction site after the poly(A) tail (SI Appendix, Supporting text). *Lingula anatina* ASIC (g20471.t1 from *Lingula anatina* Ver 2.0, OIST Marine Genomics Unit), synonymously mutated to remove internal restriction sites and including a C-terminal myc tag before the stop codon, was commercially synthesized and subcloned (Genscript) into HindIII and BamHI sites of pSP64poly(A) (SI Appendix, Supporting text). Plasmids were linearized with EcoRI, and cRNA was synthesized in vitro with SP6 Polymerase (mMessage mMachine kit, Ambion). Stage V/VI *X. laevis* oocytes, purchased from Ecocyte Bioscience (Dortmund, Germany), were injected with 3-90 ng cRNA. After injection, oocytes were incubated for one to three days at 19°C in 50% Leibowitz medium (Merck) supplemented with 0.25 mg/ml gentamicin, 1 mM L-glutamine, and 15 mM HEPES (pH 7.6). Whole cell currents were recorded from oocytes by two-electrode voltage clamp using an OC-725C amplifier (Warner Instruments) and an LIH 8+8 digitizer with Patchmaster software (HEKA), acquired at 1 kHz and filtered at 200 Hz. Currents were also analyzed in pClamp v10.7 software (Molecular Devices) and additionally filtered at 1 Hz for display in figures. Oocytes were clamped at -60 mV, unless otherwise indicated, and continuously perfused with a bath solution containing (in mM): 96 NaCl, 2 KCl, 1.8 CaCl_2_, 1 MgCl_2_, and 5 HEPES (for pH > 6.0) or 5 MES (for pH ≤ 6.0). pH was adjusted with NaOH, HCl, or KOH, as appropriate. In most experiments, activating/desensitizing pH was applied to oocytes in between resting periods (at pH 7.5 for most ASICS, at pH 9.0, 8.6, and 8.0 for *O. fusiformis, L. anatina S. californicum* ASICs, respectively, unless otherwise indicated) of at least 30 s. After retrieving current amplitude from pClamp, all data analyses were performed in Prism v9 (GraphPad Software). In concentration-response graphs, currents are normalized to maximum proton-gated current. For ion selectivity experiments, IV relationships were measured in regular bath solution and that in which extracellular NaCl was replaced with KCl. IV relationships were obtained by activating the channels at different membrane potentials from -80 to 60 mV, with 20 mV increments, unless otherwise indicated. Reversal potentials (V_rev,Na+_ and V_rev,K+_) were taken from the intersection of the IV curve with the voltage axis. These values were used to calculate relative permeability P_Na+_/P_K+_ with the Goldman-Hodgkin-Katz equation, P_Na+_/P_K+_ = exp(F(V_rev,Na+_ - V_rev,K+_)/RT), where F = Faraday constant, R = gas constant, and T = 293 K. Standard chemicals were purchased from Merck. Specialist chemicals (Fig. S1) were purchased from Santa Cruz Biotechnology (item sc-222407, sodium ursodeoxycholic acid, ≥98% purity) or synthesized by Genscript ((pyroE)WLGGRFamide, ≥97% purity – “Hydra RFamide I” from (11)).

## Acknowledgements

We thank Chris Lowe (Stanford University) for *Phoronopsis harmeri* and *Schizocardium californicum* samples.

## Notes

### Competing Interest Statement

The authors have declared no competing interest.

### Summary of Updates

Based on feedback, some discussion of gene expression was re-worked; additional control experiments were performed and data added; some figures were changed for clarity; and various text was re-worked for clarity.

