## Supporting Text and Supporting Figures 1 and 2 for "Peripheral and central employment of acid-sensing ion channels during early bilaterian evolution"

***This PDF file includes:***

Supporting text
Figure S1
Figure S2

***Other supplementary materials for this manuscript include the following:***

Dataset S1

### Supporting text. Modified pSP64 vector and ASIC inserts.

Sp6, HindIII, 5'UTR, SalI, XbaI, BamHI, Myc + TAA stop codon, 3'UTR, poly(A), EcoRI

#### Modified pSP64 Vector

ATTTAGGTGACACTATAGAATACAAGCTTGCTTGTTCTTTTTGCAGAAGCTCAGAATAAACGCTCAACTTTGGCGTCGACTCTAGAGGATCCGAGCAGAAGCT
CATCAGTGAGGAAGATCTCTAAGGTTACCACTAAACCAGCCTCAAGAACACCCGAATGGAGTCTCTAAGCTACATAATACCAACTTACACTTTACAAAATGTT
GTCCCCCAAAATGTAGCCATTCGTATCTGCTCCTAATAAAAAGAAAGTTTCTTCACATTCTAAAAAAAAAAAAAAAAAAAAAAAAAAAAAACGAATTCGTAAT
CATGTCATAGCTGTTTCCTGTGTGAAATTGTTATCCGCTCACAATTCCACACAACATACGAGCCGGAAGCATAAAGTGTAAAGCCTGGGGTGCCTAATGAGTG
AGCTAACTCACATTAATTGCGTTGCGCTCACTGCCCGCTTTCCAGTCGGGAAACCTGTCGTGCCAGCTGCATTAATGAATCGGCCAACGCGCGGGGAGAGGCG
GTTTGCGTATTGGGCGCTCTTCCGCTTCCTCGCTCACTGACTCGCTGCGCTCGGTCGTTCGGCTGCGGCGAGCGGTATCAGCTCACTCAAAGGCGGTAATACG
GTTATCCACAGAATCAGGGGATAACGCAGGAAAGAACATGTGAGCAAAAGGCCAGCAAAAGGCCAGGAACCGTAAAAAGGCCGCGTTGCTGGCGTTTTTCCAT
AGGCTCCGCCCCCCTGACGAGCATCACAAAAATCGACGCTCAAGTCAGAGGTGGCGAAACCCGACAGGACTATAAAGATACCAGGCGTTTCCCCCTGGAAGCT
CCCTCGTGCGCTCTCCTGTTCCGACCCTGCCGCTTACCGGATACCTGTCCGCCTTTCTCCCTTCGGGAAGCGTGGCGCTTTCTCATAGCTCACGCTGTAGGTA
TCTCAGTTCGGTGTAGGTCGTTCGCTCCAAGCTGGGCTGTGTGCACGAACCCCCCGTTCAGCCCGACCGCTGCGCCTTATCCGGTAACTATCGTCTTGAGTCC
AACCCGGTAAGACACGACTTATCGCCACTGGCAGCAGCCACTGGTAACAGGATTAGCAGAGCGAGGTATGTAGGCGGTGCTACAGAGTTCTTGAAGTGGTGGC
CTAACTACGGCTACACTAGAAGAACAGTATTTGGTATCTGCGCTCTGCTGAAGCCAGTTACCTTCGGAAAAAGAGTTGGTAGCTCTTGATCCGGCAAACAAAC
CACCGCTGGTAGCGGTGGTTTTTTTGTTTGCAAGCAGCAGATTACGCGCAGAAAAAAAGGATCTCAAGAAGATCCTTTGATCTTTTCTACGGGGTCTGACGCT
CAGTGGAACGAAAACTCACGTTAAGGGATTTTGGTCATGAGATTATCAAAAAGGATCTTCACCTAGATCCTTTTAAATTAAAAATGAAGTTTTAAATCAATCT
AAAGTATATATGAGTAAACTTGGTCTGACAGTTACCAATGCTTAATCAGTGAGGCACCTATCTCAGCGATCTGTCTATTTCGTTCATCCATAGTTGCCTGACT
CCCCGTCGTGTAGATAACTACGATACGGGAGGGCTTACCATCTGGCCCCAGTGCTGCAATGATACCGCGAGACCCACGCTCACCGGCTCCAGATTTATCAGCA
ATAAACCAGCCAGCCGGAAGGGCCGAGCGCAGAAGTGGTCCTGCAACTTTATCCGCCTCCATCCAGTCTATTAATTGTTGCCGGGAAGCTAGAGTAAGTAGTT
CGCCAGTTAATAGTTTGCGCAACGTTGTTGCCATTGCTACAGGCATCGTGGTGTCACGCTCGTCGTTTGGTATGGCTTCATTCAGCTCCGGTTCCCAACGATC
AAGGCGAGTTACATGATCCCCCATGTTGTGCAAAAAAGCGGTTAGCTCCTTCGGTCCTCCGATCGTTGTCAGAAGTAAGTTGGCCGCAGTGTTATCACTCATG
GTTATGGCAGCACTGCATAATTCTCTTACTGTCATGCCATCCGTAAGATGCTTTTCTGTGACTGGTGAGTACTCAACCAAGTCATTCTGAGAATAGTGTATGC
GGCGACCGAGTTGCTCTTGCCCGGCGTCAATACGGGATAATACCGCGCCACATAGCAGAACTTTAAAAGTGCTCATCATTGGAAAACGTTCTTCGGGGCGAAA
ACTCTCAAGGATCTTACCGCTGTTGAGATCCAGTTCGATGTAACCCACTCGTGCACCCAACTGATCTTCAGCATCTTTTACTTTCACCAGCGTTTCTGGGTGA
GCAAAAACAGGAAGGCAAAATGCCGCAAAAAAGGGAATAAGGGCGACACGGAAATGTTGAATACTCATACTCTTCCTTTTTCAATATTATTGAAGCATTTATC
AGGGTTATTGTCTCATGAGCGGATACATATTTGAATGTATTTAGAAAAATAAACAAATAGGGGTTCCGCGCACATTTCCCCGAAAAGTGCCACCTGACGTCTA
AGAAACCATTATTATCATGACATTAACCTATAAAAATAGGCGTATCACGAGGCCCTTTCGTCTCGCGCGTTTCGGTGATGACGGTGAAAACCTCTGACACATG
CAGCTCCCGGAGACGGTCACAGCTTGTCTGTAAGCGGATGCCGGGAGCAGACAAGCCCGTCAGGGCGCGTCAGCGGGTGTTGGCGGGTGTCGGGGCTGGCTTA
ACTATGCGGCATCAGAGCAGATTGTACTGAGAGTGCACCATTCGACGCTCTCCCTTATGCGACTCCTGCATTAGGAAGCAGCCCAGTAGTAGGTTGAGGCCGT
TGAGCACCGCCGCCGCAAGGAATGGTGCATGCAAGGAGATGGCGCCCAACAGTCCCCCGGCCACGGGGCCTGCCACCATACCCACGCCGAAACAAGCGCTCAT
GAGCCCGAAGTGGCGAGCCCGATCTTCCCCATCGGTGATGTCGGCGATATAGGCGCCAGCAACCGCACCTGTGGCGCCGGTGATGCCGGCCACGATGCGTCCG
GCGTAGAGGATCTGGCTAGCGATGACCCTGCTGATTGGTTCGCTGACCATTTCCGGGTGCGGGACGGCGTTACCAGAAACTCAGAAGGTTCGTCCAACCAAAC
CGACTCTGACGGCAGTTTACGAGAGAGATGATAGGGTCTGCTTCAGTAAGCCAGATGCTACACAATTAGGCTTGTACATATTGTCGTTAGAACGCGGCTACAA
TTAATACATAACCTTATGTATCATACACATACG

#### ASIC inserts for expression in *Xenopus laevis* oocytes

>IpulchraASIC
GTCGACATGTTACCAAGCAATCAATGGGCCATCACTGTGAATGCGACGCCAGCCCACCGCGTTGGGTCCGAGTATGATCCGCAGTACAACCCGCAGACCGCAC
GGGGGCCGGAAACCCAGGTGGTGCGGTATGCGGCGCCGGTGCGGACCGTGCTGGTGGACAGCTACTGCCAGACCAGCGCTCCGCCCTCCGCCAGCTGTCTCCC
CATTGCGGAGCGGTACCAGTGCGCCATGGCCAGTGTCCAGTACAGCGAGGACGAACACTCCATAGGAATCAAGACATTCCTATCCGAGGAAACCACCATGCAC
GGCATCAAATATGTGGTGCTGCCGGGTGCGCGGATCACGATCAAGCGGGTGCTGTGGACGCTGGTGTTCGCCCTCTTCTGGGTCTACTTCATCTCGGTCTCCG
TCAAGAAGGTCCAGTACTACTACTCCTTCCAGCACATCACCCAGGTAGACGAGGAGTCCAAGGAACTCAACTTCCCGGCCATCACGATATGCAACCTGAACCC
GATCCGGCTGTCGGCGTTCACCAAGAGCGACCTGTACCAGGCGGGCCGCATGTACCAGATCCTGTCGGCCAACCTCACCCTGAAGGAGGACCTGTTCTCCTCG
GGCGACGAGACCGCGGTCGCCCTCCGCTCCAAGACCGACTTCACCGAGCAGGACAAGCGCGTCCCCTTCGACCTCGAGGAGTTCCAGCTGCGTGCCGGCCACA
AGATGAAGGACATGCTCAAGGCGTGCAAGTGGCGCGACCAGCAGTGCACCGAGAAGGACTTCCAGCCCGTGCTGACGGAAATGGGGCTGTGCTATCAGTTCAA
CAGCAACAAGTCGCTCGACCCGGAGACGGGCGAGTACGCGTTCCGCAAGGTGTACAAGGGCGGCGCCGGCAACGGACTCAAGCTACTCCTTGTCACCCAGGAG
CAGGACTACCTTCCGCCCAACATCGAGGACTTCGACTCGACGCTGGAGACGGGGTTCAAAATCGAGGTGCACGACCCGGACGTGCCGCCGCTGGTGCACCAGC
TGGGCATCGGGCTGGCGCCCGGGCTGCAGACGTTCGTGGTGCTGCAGAAGGAGGAGGTGAACTACCTGTCCGAGCCCTGGGGCTCCTGCATCGACCCCATGGA
GCCGGGCAAGGTCACCAACTTCACCAAGTACTCGGACCCCGCGTGCCGCATCGAGTGCGAGACCGAGCACATCGTGCAGGAGTGCGAGTGCCGCATGCCCTTC
ATGCCCATTTTGCCGGCGTACGAGAACTCGAAGACGGTGTGTTCACCCGCGCAGATCATCAAGTGTGCGCGGTCCGCCCTGGCGCGGATAGAGGACGGAGAGG
CGTGCGGGCCGGAGACGTGCATCAAGAGCTGCCGCATCTCCCGCTACACCAGCTCCCTCTCCTCCGTCAAGTTCCCCAGCTTCCTCGGGGCCATCAAGCTGCA
GGAGGAGTTCCCCGACACCTTCCCCACCAAGAAGGACGTCACCCCCGCCAATGAGTCATCCGCGGCCGCCGCCTCCGCTCCCGTCACCGGCAAATGGAAAGAG
ACCATAGACGGAGCTGCGGCCCAGAAGAAATGGGAGCAGGCGCTGGTCGTGTCCATATACTTCCAGTCGCTGAGCTTGCAGAAGATCGAGCAGGTCGAGGCGT
ATGACTTCCCGGCCATGCTCGGCGACATCGGAGGTCTCATGGGGCTGTTTATCGGCGCCTCCGCCATGACGCTGTTCGAGTTTCTCTTCTTCTTCTGTGACTC
TCTCATCCTCCGGCGCAAGAGCACCAAGCAGCCGCCGCCGGACACCTCGAGCCAGAGCTCGCCCTCCAGCGGCCAGAGGAAGGTTGTCCACGCCGAGATGGAC
AACGGCAAGTTCTCCATGCCGCCCGGCGCCGTCGATGGCTCCAAATTCGTCATCAACTTCAACCCGGACTACAATCCGGGCATTACACGAGTGCTTCCGCAAT
TCACTGCTCATCCTCTGCACAACAGCTTCCAGCAGCAACAACAGCCGCAGCAGCAGCCGCGCGGCTACTCACCTCTGATGAAGCGGGCCGCCGGCAACTTCGG
AGCAACCGACGGCGGCGATGAGTCATTCGGCCAGCAGCCCGCGCCCGCCCAGCAGCCGCAGTTCAACACTCGAATGTACCAGCTGCCCAACAACCAGCCACAG
GAGCGCTACTGCGGATCC
>HmiamiaASIC
GTCGACATGAATAGGCCAATGAAATACGTCGCCACTCAACCCGTTGATCAACCCCCTCAATGCACATTCATTGATTCATATGCCGATCATGATCGAAAATCTC
TCCTCGATTTTGCCGATGAAACAACAATGCATGGAATGCGTTATGTGGTCTTGCCGGGCGCGAAAATTACCATCAAACGCGTGCTATGGACTTTTGTTTTTGC
GCTTTTTTGGGTGTACTTTATTTCAGTTAGTATTAAAAAGGTCCAATCTTATAGGAAATACCAGCATATCACCCAAGTTGACGAAGTATCCAATTCCCTTCCT
TTTCCCACTATCACAATTTGTAATTTAAACCCAGTACGGATATTTTCTTTCACAAAAAATGATCTTTATCATGCCGGTCAGATGTATGAACTACTAGATGAGA
ACATGACGTTGATCAAAGGACTCTCCAAGAATGTTAGTAAATTTCTTTCAAAACTGGCCAACTTCACAAATTTCACACCGACGAGCTTCGATCTGGGGGAGTT
TCAGCGCCGTGCAGGACATAAAATTGAAGACATGCTAATAAAGTGCAAATGGAGAGACGAAAAGTGTTCAGCAAAAGACTTCACACCCATATTGACCGAGATG
GGCCAGTGTTATCAGTTTAACAAAAACCCCATCACAAATGATAGTGGTATTCCAATTTACAGAAATGTTTCAAAGGGTGGTCAAGGCAATGGCCTTAAACTCC

TTCTTGTCACGGAAGAAGAACAATACTTGCCTCCTAACATTGATAAATTCGATGTGAGTCTTGAATCAGGATTCAAAGCTTTGGTTCACGACAACGACATTCC
TCCTTTAGTTCAGCAGTTAGGCATCGGTCTAGCTTCGGGTTTACAGACATTCGTTGTACTTCAGAAAGCAGAGGTTAAATACTTGGGCCAGCCATGGGGCGTC
TGCTTGAAATCAGAAAATTCCCCTGACAAGATTTGGAGCCGCTACAGCGATTCATCTTGTAGAATAACATGCGAAACGCAACATATTGTAAAGAAATGCAAAT
GTAGATTTCCATTTATGCCAATCTTAAAAACTGATCCTAATTCAAATACCATCTGCACAACAAAGCAAATCAAGGACTGTGGCACAAGGGAACTCGAAAACAT
CGAGGACGGTTCGGCGTGCGGACCCGATTTCTGCATCAAACCTTGTGAAATTACTCGCTACAGAAGTTCTTTGTCATCGGTAAAATTTCCCAGCGCTCAAGGA
GAAATCAAACTAAGCGACGCACTGAAGAAGCACACAGAGGGAAAAGGACGTAATTACACATCAGGTTCAGAAAAACTATCAGATCGCGCTCTTGTCGTTTCCA
TCTACTTCCAATCACTCAGTCTGGAGAAAATTGAACAAGTTGAAGCATATGACTTTGCTGCCATGCTTGGTGACATCGGAGGTCTGATGGGTCTTTTCATCGG
AGCAAGCGCAATGACATTATTTGAGTTCTTCTTCTTCTTTTGGGACTTTATTCTTCTTAGAAAAAAACCTAGCAACCCAGAATCCAATAACCGCCAGCCTCAA
ATGGCCAACGACCGGGACTGCATGCCACCGGGCGCCGTGGATGCCTCCAAATTCGTTCTTCCATTGCATAATGAATACAACCCACAGTTAATCCATGTCTCCG
CCCTTCAACATCAAAACTCTGTTCAGCTGCCAATTCCAAGAATAGAGCAATACAACAAATTTTCACCCCTCGTCAAACGCAGCGTCATGCCAGTCCATCCCTA
CTCAGTAAATTTCTTCGAAGAGAAGAGTTCGAAACTGCTTATATCCAACCCCTTTCAAGAGAAGTACTGCGGATCC
>CmacropygaASIC
GTCGACATGCATGGAGCAAGATATGTGGTGCTGCCAGGTGCAAGGATAACGATAAAGCGAGTGTTGTGGACACTGGTGTTTGCCCTCTTCTGGGTCTACTTCA
TCAGTGTGAGTGTGAAGAAAGTGCAGTTCTACTACTCCTACCAACACATCACACAGCTAGACGAGGAGTCACAGGCCTTAGACTTTCCTGCCATCACGATCTG
TAACTTAAACCCAATTCGCCTGAGTGCATTCACTGCCTCCGACCTCTACAACGCTGGCAAAATGTACGACATTCTGGATGACGACTCCAAACTGAAAGAGGAC
CTTTTTTCAGAAGACAACCCAAAGACAAAACTCCTCATGAGGACTCTGGCTTCAATGACAGATGATGACAAGAAGAAAGACTTCAACCTGGAAGAGTTCCAAA
TGCGAGCAGGTCACAAGATGAAAGACATGCTGAAAGCGTGCAAGTGGAGAGACATCAAGTGCACTTCGGACGATTTCACCCCGGTTCTGACCGAGATGGGGCT
GTGCTACCAGTACAACGTGAACAAAACATTCGACGATGAAACAGGTGACTACCAATTTAAGAAAGTGTACAAAGGAGGTGCTGGAAATGGTCTCAAAATGCTT
CTCATTACCCAAGAAGACGAGTATTTGCCACCACAGATTGAGAGCTTCGAGACCTCCCTTGAGACTGGCTTCAAGATAGAGATCCATGATCCAGACGTGCCCC
CTCTGGTCCACCAGCTGGGCATTGGCATTGCCCCTGGGCTTCAGACATTTGTGGTTCTGCAAAAGGAGCAGGTGACGTACCTGACTAAGCCGTGGGGTGAGTG
TGTGCAAAAGTACGAGAAGACTATCAACTTCACAAAGTACTCGGATCCAGCATGTAGGATAGAGTGCGAGACCCAGCATATTGTGGACGAGTGCTCCTGCCGA
TTACCCTTCATGCCAATTCTGAGCAACAAGCCCAAAAGTAGCATTGTGTGTTCTCCTCAACTGTATAAGGAATGTGCGAGAGAGAAGTTGAACAAAATAGAAG
ACGGGGAGGCTTGTGGCCCAGAGACCTGCATCAAAGCCTGCGAGATTTCAAGGTACAAAAGCAGCATGTCATCAGTGAAGTTCCCCTCTTACCTGGGTGCAAT
CAAACTGAGCGAAGAATACCCAGGCGTGTTCACAAAGGACAGCACCAACTCTGGAGGTGGGGGTGACTCGGGTGGCGACGGAGGGGGCGATTCTCGAAGTGGT
GGAGACGGAGGTTCGGATCGGTCATTTGACGGAGGGTCTGTTAAAGATGATGTGAAAGCTAAGGCTACTCAGTCTTATTGGAAAGAGGTTGAAAACAAAATAT
CAAAGCAAGCCTTGGTCGTGTCTATCTACTTCCAGTCTCTGAGTTTGCAGACTATTGTCCAGGTCGGAGCATATGACTTTGCAGCCATGCTTGGTGACATTGG
TGGTCTGATGGGTCTATTCATCGGAGCTTCTGCCATGACACTTTTCGAGTTCTTCTTCTTCTTCTGCGACTCCCTCATCCTGCGTAGGAAGGGTGCCGGCAAC
AGGAACAGCAATGCAGGAGGGGGATCCGATGATGAGAAATCAGAAATGATAGGGGACAATTTGTTGCGAGGCCACGATGGTCAACCCCTTAAGAGTGTGCTGT
TGGCTGCCCCTGGGGCCGTGGATGGGTCCAAGTTCGTGTTCAATGTCAACCACAACAACGACCACAACGCCGCCGCTGCCGCCGACTACAATGCTGCCATGTC
CAGATTACTTCCACAGTTTAGCGCCCATCCTTTGCACAACTCATTTCAACAGCAACAACAGCAACAGCAACTGTTAAACCAACAGCGAGGCTACTCACCCCTG
ATGAAAAGGGCTAACTACCCCTCGTCAGGTGCATTCGGCCAACCGCCAAACCAGAACCAACCCAACTCAAATGATGTCGGTAGTCTCTTGTCCCCCCACGATC
CCTCCAGCTACCCTGCGTCTCCAGTACCCCCCGCGAATGGTGGAATAGGGCTTCCGGTGACCTCATTTCCCGGAGCTCAGCAACAACAGAATATTCCACAGAA
CATGCTCGTCACTGCACCCTATCAACAGGGGGGTTTGCCCGGAGCGGGGGGCAAAATATTCCTGGCCAACAGTCTTCCTCAAGAGCGATATTGCTCTAGAGGA
TCC
>TtransversaASIC
GTCGACATGTCTTCTACAACACACGAACCAACAGACTCGGAGATGAGGCACAGGACTCCTGCAAATAATACACCAGGAGTTCTATCACATAATCAGCTTGAAC
TTAAAACAAAGGAAAGAATAGAAACATTCGGTGAAGATTGTGATTTCCATGGCATCAAAAGAATATTACGACATGAATATCACTTATGGAGAAGGATTACATG
GCTCATGTTTTTTGTCTTTGGTTTAACATTTTTGACATATCAAATCACAGAGTCAGTCTTGACTTACCTGAAGTATGAACATGTTACTAAAGTGGATATGATG
TACTCCAATACAATGGAATTTCCTGCCGTTACCATATGCAATCTTAACCAATTTAAAGTCTCTGCATTAACAGCTCTGGATTTGCATCACTTCGGACAAAAGC
TAGGGATTTTCCTTAATGGCACATACACCCTTACACATCCCGACCAGTACAACAGTACATGGGTACAGTGGGTGAATGGAATCAACTGGACTGAAATAGCCAG
TGGTCCTGATGACTTTGATGTGGAAGAGTTCTTTAACAGAACAGGTCATCAGAAGGAACAAATGATTTTGTTATGTCTTTGGAGAGGGGCAGCCTACAACCAC
TCTGATTTCCAAATGACACACACCCATCTAGGAAATTGTTTTACATTTAACCACGGAAAAGATGATGTTAACTTTCACACAAGAAATGCTGGAACTGAAGCAG
GTCTAAAGTTGTACTTAAATGTTGAACAAGATGAATATCTAGAAGGTGAAGACTCTGCTGATGCTGGATTCAAACTGTTAGTTCACGATCAAAAAGATCCCCC
ATTTGTGAAAGAGTTGGGCTTTGGGGTTGCTCCTGGATATCATTATTTTGTGTCTTTACAGAAACAAAAGATTAAAAATCTGTCTAAGCCATGGGGTAATTGC
CAGTCAGGACAACTGGAATACTATGACCACTACACAATTCCAGGCTGTCGGATAGAATGTGAACTGAAAACAGTCAAAGAAACATGTGGATGTCGTCTTATCG
AGATGCCAGGAAATGATACAGTGTGTCACGGCAATGCATACATGGGATGTGCATATCCTGCATTAGAGAGAGTGGAACACTCGGACAATTGTGTTTGTCAAAA
TCCATGTGAATTTACACACTATAAACATGCCACAACCTTTGTGGAGCTAAGAGAAAATACAGTGGACAGGATTCAAAAGAAAAATATTCAGTATACAAAAGAA
AAATTGAAGTCCGACCTGGTAATACTGAGTGTATACTTTGAGAGACTCAATTATGAAGAATATGAACAACTGCCAGCCTACACTCTGGTTGATCTATTCAGTG
CCATTGGTGGTAATATGGGTCTCTTTATTGGAGCAAGTGCCTTGACAGTTCTCCATTTGATGGAGTTCATTGGGTCAGAGTTAACTCTTTGGATGTGGAACAG
AAAAAACAAGAAACAAAACCAAGTAATGACAACAGTCACTCCCATGAAACTACCAATACAGGATGGATCC
>Lanatina_ASIC
AAGCTTCCTTGGCAAGCATGCAGAAAAAGACCTCGTGGATAGAGACTGCCAATCGGTTAGTCGGCAACACCAAAAGACAGAAGCAGTCAGAATCCACAGAATC
AAAAGCTAGCAAAGAAAGCGGCTCCGTCAGCGAAGGCCTGGAACAATTTGGAGAAGACACCGACTTCCACGGACTGAAGCGTGTCTTCAGGAGAGACTATACA
CTGTTACGCAGAACCTTTTGGTTGTTTTTCTTTCTGGCGGGTCTTACGGGTTTTATCGCCAACACAGTTGACAGGCTGAGCTATTACTTGCAGTATCCACACT
CGTCTGAGCTGGACGTGATGTACGATATGGAGTTAGAGTTTCCAGCCGTAACCATATGCAATATGAACTCCTATAGGCTATCCGCTTTGACAGACGTTGATAT
ATTACATTTTGGTGAAAAACTACACATTTTAGACGAAAACCGCCGTCTTATACACCCGGAATATTATAACCAAAGCTGGGTGGATTGGGTGCACGGTATTAAT
TGGACAGAGTTAGCTGCCAACGATGACGAAGAAAATTTTGACGTGCTTGAATTTATACGTCGAACCGGCCACCAGCTGGAAGACATGATACTTCTGTGTTCGT
GGAAAGGGCAACACTGTGGCCCGGAAAACTTTACCTCGGTTTTCACTCATTTTGGCATCTGTTATACTTTCAATGCTGACCACTCTTATAAATACGTATCACG
AAAGGCGGGGGCGGGGAACGGGTTGAAGTTATATATCAACATCGAAGAAGAGGAGTATTTGACTTCGGATGTCCTAGCAGGGCAAGATGCTGGACTGAAGATG
GTTATTCACGCTCAGGAAGAGCCGCCGTTTGTAAAGGAGTTGGGATTTGGAGTGCTACCGGGGGACCACCACTTTATAGCCATACAGAAGAAATACGTTCACA
ACTTGGTTCCTCCATGGGGAAATTGCTACGACGGGAAACTCAAGTATTACTCCCATTATAGCGTCCCTGCCTGTAGGATAGAGTGTGAGACAGATACCATCGT
AAAAGAGTGTGGCTGCAAGCTGGCTGAAATGCCAGGCAACACATCCGTATGTCTAGGACATATGTACATGGGATGTGCCTATCCAGCCTTAGAGGAAGTGGAG
CACTCAGACCTGTGCTCCTGTCAGAATCCATGTGATATGACCACCTATAGGCGAGACATCTCTTCCGTAAGGCTTCGAGATACTACCTTAGATATCATAGCCG
AAAATAACCCGCAAGTTAAACGGGAAACACTAAGAGATAACCTACTGGTGTTGAACATCTATTACGAAGAGCTGTGCTATGAGACCATCAGACAAATCAAAGC
ATATTCCATACCAGCATTACTTAGCGACATAGGAGGCCAGATGGGTTTGTTTATAGGAGCTAGTGTTCTGACTCTGCTGCATGTGATCGAGGCCGTTGGGGCT
GTTGTTGGTGGAAAGTTCCTCAAAACAACGAAGCAAGGGCGTGTCAGTACAACTCGGGTCCAAAGTTTGAAGGGATCCGAGCAGAAGCTCATCAGTGAGGAAG
ATCTCTAA
>PharmeriASIC
GTCGACATGGAAGAGCCGGAGACAGAACCTCTCACACAGAACGGACTGAACCAACGTTTGGGAAAGAAAGCGGTCCAGGACTGTATTATAATTGACTTGGTCA
ACAATGAAATACCTGAGCACGAGCGAAAGGAAACGTTGAAGACTCGTTTTGAAAACTTTGGAGAAGACACCGACGTCCACGGATTAAAATACATACTACGCGA
GGACAGTCCAACGTATAAAAAAGTTATATGGATGATGTTGCTTGTAATTAGCCTTGGCTACATGACATTTGAAATTTATGGAAGGTTTTCGTGGTATTTTACT
TACCCACATGTCACCAAGGTCGATGTTGTGTTTCGCAATAACATGGAATTTCCTGCATTCACCATTTGTAATATGAACAAATTCCGCGCGTCGGCCATGACAG
ATGTGGATATATTAAACATGGGAAAAATACTAGGAATTGTGAATGACAATATGAAACTGCACCACGGTGATCATTATAACGAAACATTCGTGAACTGGGTAAA
CAGCATGAACTGGACACGCGTCCGAGAGAAAAACAAAACATTTCACCTGGAAGAATTTTCTAGACGAGTTGGGCATCAGGCCAAAGACATGATTGTCTACTGC

CGATGGAAGGATGAGTTATGCGGACCTGACAATTTTACTCACAGCTTCACGCATCTTGGAAACTGCTATACGTTTAACGACAATCAAAAGTTTGCGGCCCGAA
AGGCGGGTGCCGGCAATGGACTAAAGTTATATCTTGACGTTGAAGAATTTGACTATCTTGAAACTGCAGATGCCGCAGACGATGGCCTGAAGGTCATAGTTCA
CAGTCAAAAAGAACCGCCTTTTATAAGAGAACTTGGATTTGGACTTATGCCGGCACAACACCATTACATTGGTATAAGGAAAGCCGAGATAATCAATTTACCA
AAACCATATGGGACCTGTGCAGAAGATTTTCCAATGCGAATGTTTGAGCATTATACAATACCTGGATGTAGAATCCAGTGTGAAACGGAACACGTTGTACAGG
CTTGTGGATGTCGTCTGCCAGAAATGCCGGTGTCATCTGACGCACCAATATGTTCGCCTCTTCAGTACGACGACTGCGCACTGGCAGAATTAATACGTGTCAG
CGAGTCAGACGACTGTGTATGCCAAAGCCCGTGCCACCTCGATGACTTCCGGCTAACACATTCCAGTGTTAAACTACGCCCAGAAACAATAGAAAAGCTTCAG
ACCTTGCACGCACACGTTCCGAAACATCTCAAAGCGCATAATATTGTTGTGATGGACGTCTTCTTCGAGGCGTTGAGTCTAGAACTTATAGAGCAAAAAGTGG
CGTACCCGTGGCCAAGCTTACTTGGCGACATTGGCGGGCAGATGGGCTTATTCATTGGCGCCAGCGTCCTGACAATCCTGCATGCTGTCGAGTTTTTCACCGA
TGAAATCGCTAAGAGTTGTAAAAAGAAAGCAGACAAGACGGACCAAGTGAACCCAGATGAAACTAGGGCGTCCATGGGTCCCAACGAATCTGCCGTGGTCGTC
AAGGAAGCTGTGATAGGATCC
>OfusiformisASIC
GTCGACATGACTGAAACAAAGGAAAAACAAACTGATCCTGTCATTGAGCATTCATTTTCAACAAGACTTAAGGAATTTGGTGAAGATGTTGAATTTCATGGAG
TGAGACATCTGTTTAGAGATAATTCAATTGTGACAAAAGTGACATGGATTTTACTGTTTCTGTGTGGATGCAGTTGGCTGACCTACCAGATACATGATCGAAT
TATATATTATTATAAATATCCGCATATAACCAAAATTGATAAACTTTATGTTCCTTCATTAGATTTCCCAACAATCACAATTTGCAACATCAATACATTCAGA
AGGCACAAATTAAATGAGTATGACCTCCTTCACTATGGAACAAGCCTGAATATTCTTGATGAGAACCGGACCCTTCTTCACCCAGAGCACTATGATAAAGCCT
TTGATGACTGGGTTTACAGTATCAATTGGACAGATGTGGAGATCCATGACCGTTCCATCAATCACAGCATGGAGGAGATGTATGAAAGAGCAGGACACCAGAT
AGAGGATATGCTCATCTACTGCAAATGGAAACAACAGGAGTGTTCTGTTGCTAACTTTACCTTGATTAACACTCACTATGGGCGATGCTATCAATTCAATTCT
GGCAAGGATGGTGTCAAACATCAGTCATTTAAAGGAGGTAAAGCAAATGGTTTGAAACTTTATCTAAATGTAGAGGAACTGGAGTACCTAAACTACATGGAAG
CTTCAGACCTTGGCTTTAAAATTCTGGCACATGACCAGGATGAACCACCACTTATACAGGAATTGGGATTTGGAGTGACTACTGGCAACCACTACTTTATTGC
ATTGGAAACAGAAAGGGTGACAAGCTTACCAGACCCTTATGGCAACTGCGAAGAGGACCACAAACTTGACCATTATGATCATTACAGCATTCCTGCGTGCAGA
ATTGAATGCGAGACATTAATAGTTGAGGAGAAATGTTCATGTAGGCTTGTTGAGATGCATGGTGATCATGGTATCCGTGTTTGTACTGCTGAGGAATATCATG
ACTGTGCTCTGCCAACATTAGAGTCGATTACTGAGAGTGACACATGTGTATGTCAGAACCCATGTGAGCTGACCCAGTTTCAACACTCTATATCCTCTGTAAA
GCTTCGTGAGAGTGCAGTTGAGATGATACATGGCCACACTTCAAGCAACATAAATCTGACTGAGTTTAAGTCAGTGGAGTTCATTGAGAAAAACCTTCTTGTT
GTGAACCTCTTCTTTGATTCATTGAATTACCAGTACATTGAACAGACGGTTGCATATCCTGGTGTGTCACTATTAAGTGATGTTGGAGGTCAGATGGGATTGT
GCATTGGGGCCAGCATTCTGACAGTTTTACATTTGGTTCAGTTTGCAATTGGGGAAATCATTAAGAAGTTTCAGAAGAGAGAGAATAAAGAAGTCAACACCAA
TGTTATTCATGTGGCATCTGCAAATCCTGTAGATGATAAACTTGGATCC
>Scalifornicum_ASIC
GTCGACATGGAGCTACGGGACAACGGTTACGGGATCGGTCGGTCCCGTAAAATTAACAACAATGCTTACTCCGAATCTCTATCGTTTAACATCGCTAGTCGCT
CCTTGTCAACGGACGGTTCGAACTTAGAAAGCAATTCCCGCTCGGCGTCTTTACGGTGGGTGCATTGGGCATCGCAGGTGTCTGATATACACGGAGTTAAACA
CATTGTCAGCGAATCTTCACGTTTTCGTAAGCTGCTTTGGACGCTCTTTGTGTTAACTTCATTAGGCGTGCTTCTTTTCCAGTTTTGTGAGTCAGCATGGAAC
TATGCTCAATTCTACCACATCACTAAACTTGACGTTGAATATCTACCTCATATGCCATTTCCCGCCGTTACTGTTTGCAATTTCAACAAATACAGAAGATCGG
CGATCACACCTACAGATATGGTCCACATTGGTCAACCATTAGGTCTGGTGGACACGGAGAGAAATTTAAACAATCCAGAGTTATTCAGTGATGAGTTTATGGA
CAAATGGAACAATACTGATTGGGAAGTAGAAATGAAAAAGCCGTTCGACTTCACGGAGTTCACACGCAGAGCAGGACATCATTTTGACGAGACCATCCTCGAG
GCTCTTTGGAATGGTCATTCATGTACCCCAGAAGACTTCAGAGACTTTCTAACTCACTATGGCAACTGCTTCATCTTCAATCAATACGAAGAAGATGAAGACC
AATACCACTCAATGAGAGCTGGAACAGGTAATGGTTTGAGAATTGTGCTTGATGTCCACTCCCATGAGCATATACCGACGACAGACCTGGAGGATTCATTTAT
TAATGTTGGGTTTAAATTGATGATTCATTCCCCACACGAACCACCGTATCTTAAACAACTAGGATTTGCTGTTGGACCGGGAAATCATTATTTTCTAGCACTT
AAAAGACAAGAGATAATTCGATTATCAACTCCATACACTAGCAAAGTATGTGAAAAGTCTTCAGAAGGAACCAAACACTTCCATGAATACTCAATGTCAGCAT
GTCGAATCGAATGTGAGACTTCATTGCTCGTTCAAGAGTGTGGATGCAAACTTGTTGAGCAGCCTGGCAACGCGTCAGTGTGTTCACCCCAGCAGGTCGCACT
GTGTGCTCACGAGACATTAGATGAATACATTGAAGGACACATAGAATTTGACTGTCCCTGTGATATACCTTGCAAAAGTGAGTTGTACCCAGTTGATGTCTCT
AACAGTGGTTTGAAGAGGGATATCACCGGTAAATCAGTTGGATTGTCCAACTACTCAATAGAATACATCAAAAATAATATTGTCATGCTGACGATATTTTATG
AAGAACTAAACTTTGAGACGATCGAACAGCTTCCAGAAATGTCTATCGTAGACTTGCTTGGTCAGCTTGGTGGCAACATGGGGCTGTTTCTTGGTGCTAGTAT
ACTTACCATTTTCCAAATTTTCGAATATATCTTTGATGAGTTCAAATTTTGTATATCGCTTGGGGCTGATCGCAACCAACAGAAGAGAAAGAACAAGAAGAAA
ATTTATGAATGTGATGACAAAAGCCTTCAAGCACCATTGTCCATACCATCACAGCAAGCAGGCATAATACGTGGATTTAGAAATACGACTGTGGGATCC

**Figure S1. DEG/ENaC gene tree.**

Maximum likelihood trees of 700 DEG/ENaC amino acid sequences from 47 animals (using LG + G substitution model; see Materials and methods, Survey and phylogenetic analysis). Branch support inferred by fast likelihood-based methods aBayes (A and C) and aLRT SHlike (D). Aaur, *Aurelia aurita* (Cnidaria); Agra, *Acanthopleura granulata* (Mollusca); Apla, *Acanthaster planci* (Echinodermata); Aque, *Amphimedon queenslandica* (Porifera); Bbel, *Branchiostoma belcheri* (Chordata); Bpli, *Brachionus plicatilis* (Rotifera); Cele, *Caenorhabditis elegans* (Nematodea); Cgig, *Crassostrea gigas* (Mollusca); Cmac, *Convolutriloba macropyga* (Xenacoelomorpha); Crob, *Ciona robusta* (Chordata); Cscu, *Centruroides sculpturatus* (Arthropoda); Csub, *Childia submaculatum* (Xenacoelomorpha); Dgig, *Dendronephthya gigantea* (Cnidaria); Dgyr, *Dinophilus gyrociliatus* (Annelida); Dmel, *Drosophila melanogaster* (Arthropoda); Dpul, *Daphnia pulex* (Arthropoda); Esen, *Epiphanes senta* (Rotifera); Hmia, *Hofstenia miamia* (Xenacoelomorpha); Hspi, *Halicryptus spinulosus* (Priapulida); Hvul, *Hydra vulgaris* (Cnidaria); Ipul, *Isodiametra pulchra* (Xenacoelomorpha); Lana, *Lingula anatina* (Brachiopoda); Llon, *Lineus longissimus* (Nemertea); Lsqu, *Lepidodermella squamata* (Gastrotricha); Lvir, *Lineus viridis* (Nemertea); Mlei, *Mnemiopsis leidyi* (Ctenophora); Mmem, *Membranipora membranacea* (Bryozoa); Msti, *Meara stichopi* (Xenacoelomorpha); Nano, *Novocrania anomala* (Brachiopoda); Ngen, *Notospermus geniculatus* (Nemertea); Nvec, *Nematostella vectensis* (Cnidaria); Nwes, *Nemertoderma westbladi* (Xenacoelomorpha); Odio, *Oikopleura dioica* (Chordata); Ofus, *Owenia fusiformis* (Annelida); Pbac, *Pleurobranchia bachei* (Ctenophora); Pcau, *Priapulus caudatus* (Priapulida); Pfla, *Ptychodera flava* (Hemichordata); Phar, *Phoronopsis harmeri* (Phoronida); Pvit, *Prostheceraeus vittatus* (Platyhelminthes); Pvul, *Pontonema vulgare* (Nematoda); Rnor, *Rattus norvegicus* (Chordata); Scal, *Schizocardium californicum* (Hemichordata); Smed, *Schmidtea mediterranea* (Platyhelminthes); Spad, *Spadella sp* (Chaetognatha); Tadh, *Trichoplax adhaerens* (Placozoan); Ttra, *Terebratalia transversa* (Brachiopoda); Xboc, *Xenoturbella bocki* (Xenacoelomorpha); Xpro, *Xenoturbella profunda* (Xenacoelomorpha). Reponses to increased proton concentrations at oocytes expressing channels closely related to ASICs are shown: heterotrimeric cnidarian HyNaC (pink); rat BASIC (orange). Other ligands are the peptide Hydra-RFamide-I ((pE)WLGGRFamide; pE; pyroglutamate) and the bile acid sodium ursodeoxycholic acid (NaUDCA). Schematic phylogenetic tree of main animal groups (B).

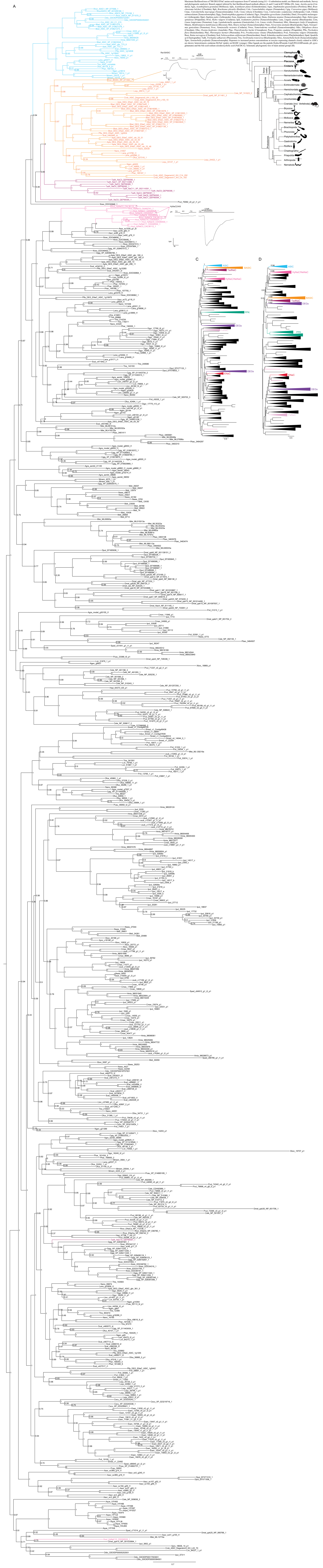

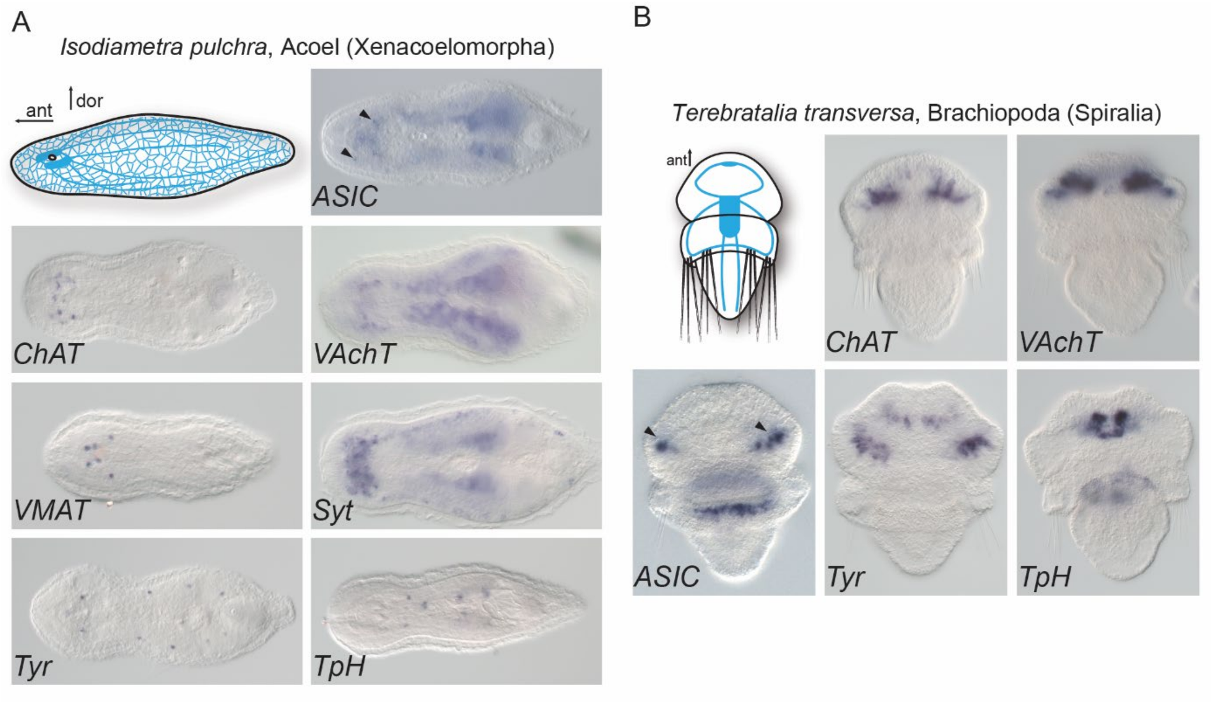


**Figure S2. Expression of neuronal markers in Xenacoelomorpha (A) and Spiralia (B).**
The neuronal markers choline acetyltransferase (ChAT), vesicular acetylcholine transporter (VAChT), vesicular monoamine transporter (VMAT), synaptotagmin (Syt), tyrosinehydroxylase (Tyr) and tryptophan hydroxylase (Tph) are mostly expressed in brain and nerve cords (A) or neuronal ganglia (B), similar to ASIC. Black arrowheads, ASIC expression. ant, anterior. dor, dorsal.
